# A cardiac microphysiological system for screening lipid nanoparticle/mRNA complexes predicts *in vivo* efficacy for heart transfection

**DOI:** 10.1101/2025.03.07.642075

**Authors:** Gabriel Neiman, Mauro W. Costa, Hesong Han, Sheng Zhao, Tammy Ng, Brian Siemons, Tomohiro Nishino, Yu Huang, Shyam Lal, Kenneth Wu, Luke Judge, Bruce Conklin, Deepak Srivastava, Niren Murthy, Kevin E. Healy

## Abstract

Lipid nanoparticle (LNP)/mRNA complexes have tremendous potential for treating cardiac diseases but lack the transfection efficiency required for successful heart therapeutics. A challenge preventing the development of cardiac LNPs is the lack of *in vitro* screening platforms that maximize transfection and predict efficacy. Here, we demonstrate that a phenotypic cardiac microphysiological system (MPS), constructed from a human induced pluripotent stem cell cardiomyocytes (hiPSC-CM) Cre-reporter line, can identify LNP/mRNA complexes that diffuse efficiently within 3D cardiac micromuscles and transfect cardiomyocytes with high efficiency. Successful LNP formulations contained an acid-degradable PEG (ADP)-lipid, where ADP-LNPs containing 10% PEG had enhanced diffusion and gene editing efficiency in the cardiac MPS. The *in vivo* delivery of LNP/mRNA complexes, including luciferase and CRE mRNA, into Ai6 mice confirmed the cardiac MPS’ screening outcomes. It revealed that ADP-LNPs achieved notably superior transfection in the heart with reduced off-target liver uptake compared to a standard LNP formulation. The cardiac MPS showed strong LNP transfection *in vitro* and pinpointed a promising formulation for *in vivo* mRNA delivery to the heart.

---

Heart failure (HF) is a global pandemic^1^ and the leading cause of death worldwide.^2^ HF can arise from a variety of sources, most often from coronary artery disease and myocardial infarction (MI).^3^ In past decades, the HF prognosis has slightly improved; however, due to sparse innovation in HF therapeutics, mortality remains high.^4, 5^ A primary reason behind the lack of progress in heart therapeutics is the inability to use phenotypic human tissue-level approaches to discover robust therapies. In recent years, there have been significant advances in the development of non-animal models (NAMs) such as organ-on-a-chip microfluidic culture devices (i.e., microphysiological systems (MPS)), which recapitulate organ-level and even organism-level functions.^6, 7^ Furthermore, since the FDA no longer requires the use of animals in master files for new drug applications, MPS are quickly becoming representative of the future of disease modeling and drug screening, therefore paving the way for complex *in vitro* models to dominate the preclinical drug discovery landscape.^7-9^

mRNA based therapeutics have great potential for treating HF, however transfecting heart tissue with mRNA is challenging since cardiomyocytes are non-proliferative and the 3D cardiac muscle is dense creating diffusional barriers posed by the extracellular matrix (ECM).^5, 10^ Recent innovations in developing non-viral vectors like lipid nanoparticles (LNPs) hold great promise for overcoming these limitations and can make a breakthrough in cardiovascular medicine due to the transient nature of mRNA transfection. Compared to viral delivery systems, LNPs exhibit advantages such as lower immunogenicity and toxicity, better cell specificity, high modifiability, and improved productivity.^11^ However, there has yet to be an effective LNP formulation for therapeutic mRNA delivery to the heart.^5^ A key challenge preventing the development of LNPs that can transfect heart tissue is the lack of *in vitro* screening platforms that predict *in vivo* efficacy. Due to the complex nature of tissue, 2D cell culture screens for LNPs have shown poor correlation with *in vivo* efficacy.^12^ Consequently, screening of LNPs for cardiac transfection can currently only be done *in vivo*, which is challenging and time consuming since current LNP barcoding approaches do not work for the adult heart.

To address the heart delivery challenges described above, we investigated LNPs with varying levels of PEGylation. Previous works showed that PEG can dramatically increase the diffusion of nanoparticles in tumor tissue; however, it can also decrease the transfection efficiency of LNPs by lowering uptake and endosomal disruption.^13, 14^ To overcome the challenges associated with developing densely PEGylated LNPs a new class of LNPs was investigated, which incorporates an acid degradable PEG-lipid (ADP) that releases its PEG chains in the endosome, and enables high levels of PEGylation on LNPs without comprising transfection efficiency.^15-17^ The newly created ADP-LNPs are built on a novel acid-degradable linker that quickly hydrolyzes in endosomes, and maintains the stability required for synthesizing LNP formulations.^18^

In this report, we demonstrate that the phenotypic tissue-level cardiac MPS **(Extended Data Fig. 1**)^19-23^ can identify LNP formulations that diffused efficiently into dense 3D cardiac tissue and efficiently transfected cardiac cells in culture, which are key requirements for *in vivo* heart tissue transfection and cannot be accurately recreated in 2D cell culture models. The tissue level cardiac MPS consists of a three-dimensional cardiac micromuscle derived from human induced pluripotent stem cells (hiPSCs) and has shown superiority to 2D monolayers of hiPSC-derived cardiomyocytes (iCMs), particularly with respect to contractility and inotropic drug response.^21-24^

Collectively, our work shows that by using the cardiac MPS as a preclinical model for *in vitro* screening of LNPs in dense 3D cardiac micromuscles, we successfully identified an ADP-LNP formulation for significant *in vivo* mRNA delivery to the heart, reducing time to discovery and animal numbers, and at significantly reduced overall costs. We foresee using this effective LNP delivery system as a starting point to develop novel therapeutics to treat HF and genetic disorders in the heart.

## Results

### PEG has high diffusivity in cardiac MPS

PEG can enhance the diffusion of nanoparticles in brain tissue due to its minimal interactions with the proteins in the ECM.^25^ However, the affinity of PEG for cardiac muscle has never been investigated and low non-specific interaction with PEG may be essential for enhancing diffusion through the heart. To determine if PEG is a suitable candidate for enhancing LNP diffusion through cardiac tissue, we measured its linear PEG diffusion through the tissue in the MPS and compared its diffusivity against the hydrophilic polymer dextran. In addition, to obtain insight into the optimal size of PEG for incorporation into the LNPs, we assessed the diffusive transport of PEG-FITC polymers with varying molecular weights (ranging from 1 to 40 kDa) within the 3D cardiac micromuscle and compared them to 4 and 70 kDa Dextran-FITC molecules. We infused PEG-FITC polymers through the media channels of the MPS, allowed the concentration within the microtissue to reach a steady-state overnight, and subsequently ran a fluorescence recovery after a photobleaching (FRAP) experiment (**Extended Data Fig. 2a, Video s1**,**s2**). We observed that the half-time to recovery was similar across the range of PEG molecular weights tested that had various hydrodynamic radii (R_h_)^26^ (1.13-6.59 nm) (**Extended Data Fig. 2b**); however, 70 kDa dextran recovery was significantly slower with a similar R_h_ to the high molecular weight PEGs (20-40 kDa), suggesting some interaction with either the cardiac MPS or the microtissue. The diffusivity of PEG-FITC polymers ranged from 85-75 μm^2^/s, with 1 and 2 kDa PEG having the highest mean values (**Extended Data Fig. 2c**). We therefore created ADP-LNPs using 1 and 2kDa ADP-PEG to promote LNP diffusion within the micromuscle.

### Development and characterization of ADP-LNPs

As demonstrated above, PEG has high diffusivity through cardiac tissue and consequently has minimal interactions with the ECM. We therefore generated LNPs with high levels of PEGylation to generate an LNP that could diffuse through tissue efficiently. Molecular weights of PEG of 1 and 2kDa were chosen for grafting onto LNPs based on their high diffusivity through cardiac tissue and their ability to efficiently generate LNPs. Conventional LNPs cannot tolerate high levels of PEGylation since it lowers cell uptake and endosomal disruption. To overcome this limitation, LNPs were formulated with a newly developed acid degradable PEG-lipid, which rapidly hydrolyzes in the acidic environment of the endosome.^18^ ADP-LNPs exhibited rapid hydrolysis in endosomes and can efficiently deliver mRNA *in vitro* and *in vivo*, despite their high levels of PEGylation **(Fig. 1a,b)**. We performed 2D and 3D cardiomyocyte screenings of ADP-LNP formulations with different PEG molecular weights (1kDa and 2kDa) and various ADP-lipid mole percentages ranging from 1% to 40% (**Table 1, 1S**). The size of the ADP-LNPs was ∼100 nm. All formulations exhibit an encapsulation efficiency (EE) above 80%, except for the 20% and 40% formulations, which have EEs of 77% and 63%, respectively. We compared their efficiency with a standard LNP formulation **(LNP-1)** made with a non-degradable 1.48% PEG-lipid conjugate, whose size was 144 nm (**Table 2**).

**Fig. 1:**
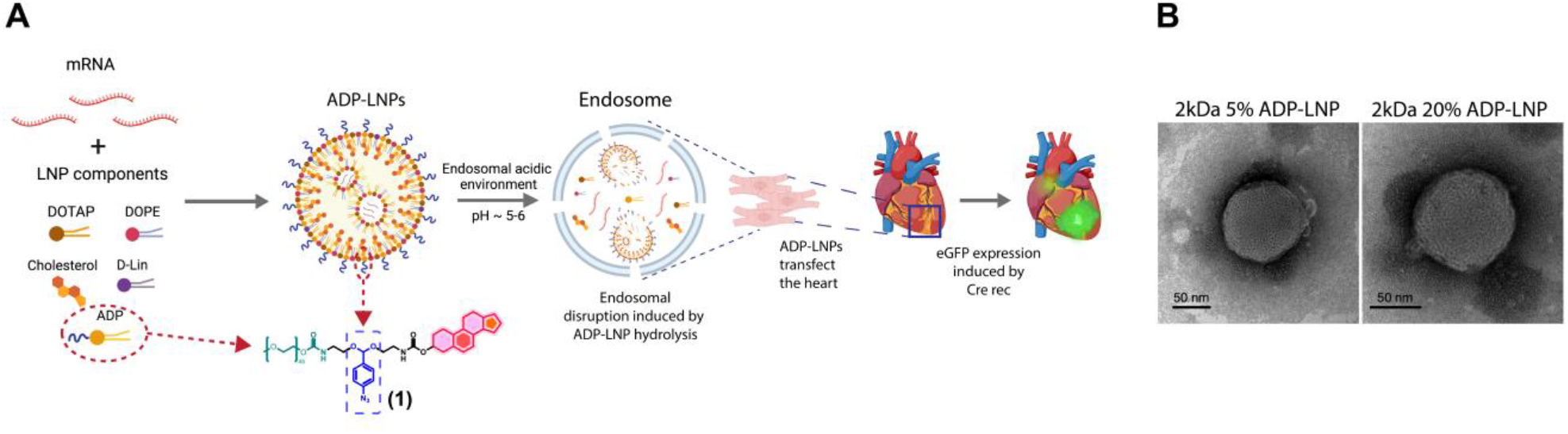
Production and characterization of acid degradable LNPs. **A**. Schematic illustration of LNP production containing an acid degradable PEG-lipid which improves the transfection efficiency and endosomal escape of LNP/mRNA complex. **B**. Transmission electron microscopy (TEM) images of LNPs with different ADP-lipid mole percentages.

**Table 1.** Components of ADP-LNPs with 2kDa PEG molecular weight.

| ADP-LNPs | Molar ratios / Molar percentages (%) |  |  |  |  | Size (nm) | %EE |
| --- | --- | --- | --- | --- | --- | --- | --- |
|  | DOTAP | Dlin-MC3-DMG | DOPE | Cholesterol | Compound 1 |  |  |
| 1% | 35/20.06 | 45/26.12 | 22/12.44 | 70/40.38 | 1.7/1 | 175.1 | 93% |
| 5% | 35/19.25 | 45/25.06 | 22/11.93 | 70/38.76 | 9.0/5 | 97.8 | 88% |
| 10% | 35/18.24 | 45/23.74 | 22/11.31 | 70/36.71 | 19.0/10 | 96.9 | 86% |
| 20% | 35/16.21 | 45/21.10 | 22/10.05 | 70/32.64 | 42.0/20 | 98.4 | 77% |
| 40% | 35/12.16 | 45/15.82 | 22/7.54 | 70/24.48 | 114/40 | 92.1 | 63% |

**Table 2.** Components of LNP-1 (DLin-MC3-DMG-LNP).

| LNP-1 | Molar ratios/Molar percentages (%) |  |  |  | Size (nm) | %EE |
| --- | --- | --- | --- | --- | --- | --- |
|  | PEG2K-DMG | Dlin-MC3-DMG | DOPE | Cholesterol |  |  |
| 38.8% | 2.5/1.48 | 62/36.69 | 40/11.93 | 65/38.76 | 144 | 92% |

### XR1 WTC hiPSC line development for gene editing efficiency quantification

To perform the ADP-LNP screening, we aimed to combine a Cre-hiPSC reporter line with our cardiac MPS as a powerful tool for the optimization of molecular therapeutics by pursuing an approach similar to the Ai6 murine reporter.^27^ Therefore, we generated a genetically modified hiPSC line (XR1 hiPSC) containing an eGFP gene downstream of a STOP cassette that is removed upon Cre delivery, making it an excellent reporter for gene editing in hiPSC-derived cells and for the novel tissue level *in vitro* approach described later **(Extended Data Fig. 3)**.

### High transfection efficiency was achieved in 2D XR1 iCM monolayers

To quantify transfection efficiency, we incubated 2D iCM monolayers overnight with the standard LNP-1 formulation and various ADP-LNP formulations containing eGFP mRNA. We obtained 80% eGFP+ iCMs when they were incubated with 1-2 kDa ADP-LNP, with mole percentages of PEG-lipid ranging from 1% to 20% **(Fig. 2a-c)**. ADP-LNPs exhibited a significantly higher transfection efficiency compared to the RNA iMAX/eGFP mRNA control **(Fig. 2c, Extended Data Fig. 4a)**. We also observed that 2kDa 40% ADP-LNPs containing eGFP mRNA were highly cytotoxic; therefore, their transfection efficiency was not quantified. Lastly, the standard LNP-1 formulation also showed an increased transfection efficiency when compared to the RNA iMAX control, but with a lower percentage of eGFP+ cells when compared to specific ADP-LNP formulations **(Fig. 2c, Extended Data Fig. 4a)**.

**Fig. 2:**
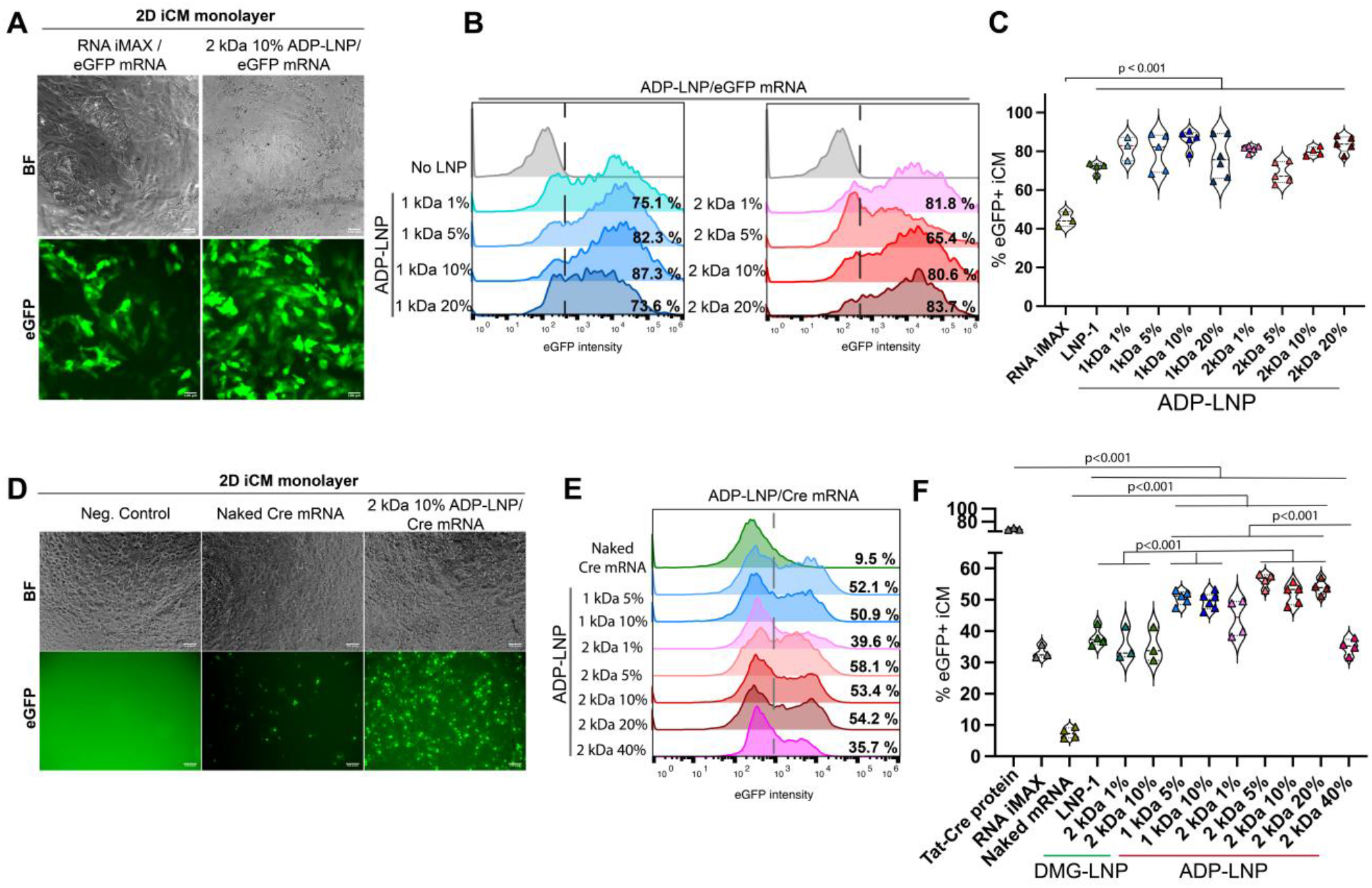
ADP-LNPs can transfect human iCMs in 2D. **A**. Brightfield and fluorescent microscopy images of 2D XR1-iCMs exposed overnight to different LNP formulations containing eGFP mRNA. **B**. Flow cytometry analysis of 2D XR1 iCM monolayers transfected with LNPs/eGFP mRNA. **C**. Quantitative analysis of B (n=3-5). **D**. Brightfield and fluorescent microscopy images of 2D XR1-iCMs exposed overnight to different LNP formulations containing Cre rec mRNA. **E**. Flow cytometry analysis of 2D XR1 iCM monolayers transfected with LNPs/Cre mRNA. **F**. Quantitative analysis of E (n=3-5). Global one-way ANOVA was performed followed by post-hoc Tukey analysis.

### Robust gene editing efficiency was achieved for specific ADP-LNP formulations in 2D XR1 iCM monolayers

We exposed 2D XR1 iCMs overnight to the various LNP formulations as described in the previous section, but now containing Cre mRNA. We observed 50-55% eGFP+ iCMs when cells were incubated with ADP-LNPs whose PEG-lipid mole percentages ranged from 5% to 20% regardless of the PEG molecular weight **(Fig. 2d-f)**. These ADP-LNP formulations showed a significantly higher gene editing efficiency compared to XR1 iCM monolayers treated with either the standard LNP-1 formulation or naked Cre mRNA without a delivery vehicle **(Fig. 2f, Extended Data Fig. 4b)**. We further evaluated LNP formulations with the same composition as the 2 kDa 1% and 10% ADP-LNPs, substituting ADP with PEG2K-DMG. As anticipated, these DMG-LNP formulations demonstrated significantly lower gene editing efficiencies compared to their ADP-LNP counterparts, underscoring the critical role of ADP in enhancing the performance of LNPs. Lastly, we observed the highest gene editing efficiency for the recombinant cell-permeant Tat-Cre recombinase protein, resulting in 70% eGFP+ iCMs. Interestingly, the gene editing efficiency upon Cre delivery was considerably lower compared to the eGFP mRNA transfection efficiency, most likely due to a different mRNA encapsulation efficiency in the LNPs or to the more complex gene editing process involved after the LNP/Cre cell transfection.

### 2 kDa 10% ADP-LNP enhanced gene editing efficiency in the 3D cardiac micromuscle

Based on the 2D iCM monolayer results, we selected Cre mRNA ADP-LNP formulations with the best-performing gene editing efficiency and achieved a similar screen in the structured 3D cardiac micromuscles within the MPS. To this end, we optimized a volumetric nuclear segmentation of the entire microtissue to assess the eGFP intensity distribution in the cardiac cells and quantified the percentage of eGFP+ iCMs for every LNP formulation **(Fig. 3a-c, Video s3)**. We observed that when the 2kDa ADP-lipid mole percentage of the LNP formulations was increased from 1% to 10%, the gene editing efficiency improved significantly, resulting in ∼25% eGFP+ iCMs in the micromuscle **(Fig. 3d, Extended Data Fig. 5a, Video s4)**. Cardiac microtissues treated with 1 kDa ADP-LNPs exhibited the same trend for the LNP formulations whose ADP-Lipid ranged from 5% to 10% **(Fig. 3d)**. Interestingly, unlike what we observed in 2D iCM monolayers, 1 kDa 10% ADP-LNP exhibited a significantly lower gene editing efficiency compared to 2 kDa 10% ADP-LNP. Furthermore, Tat-Cre protein exhibited the highest gene editing efficiency in 2D cardiac monolayers but only 3% eGFP+ iCMs were found in the cardiac MPS **(Fig. 2d-f, Fig. 3d, Extended Data Fig. 5a)**. These data reinforce the requirement to conduct screening studies in complex 3D micromuscles. In line with the previous results, the standard LNP-1, which contains 1.5% non-degradable PEG-lipid, showed a significantly lower gene editing efficiency compared to 1-2kDa ADP-LNP with high PEG mole ratios.

**Fig. 3:**
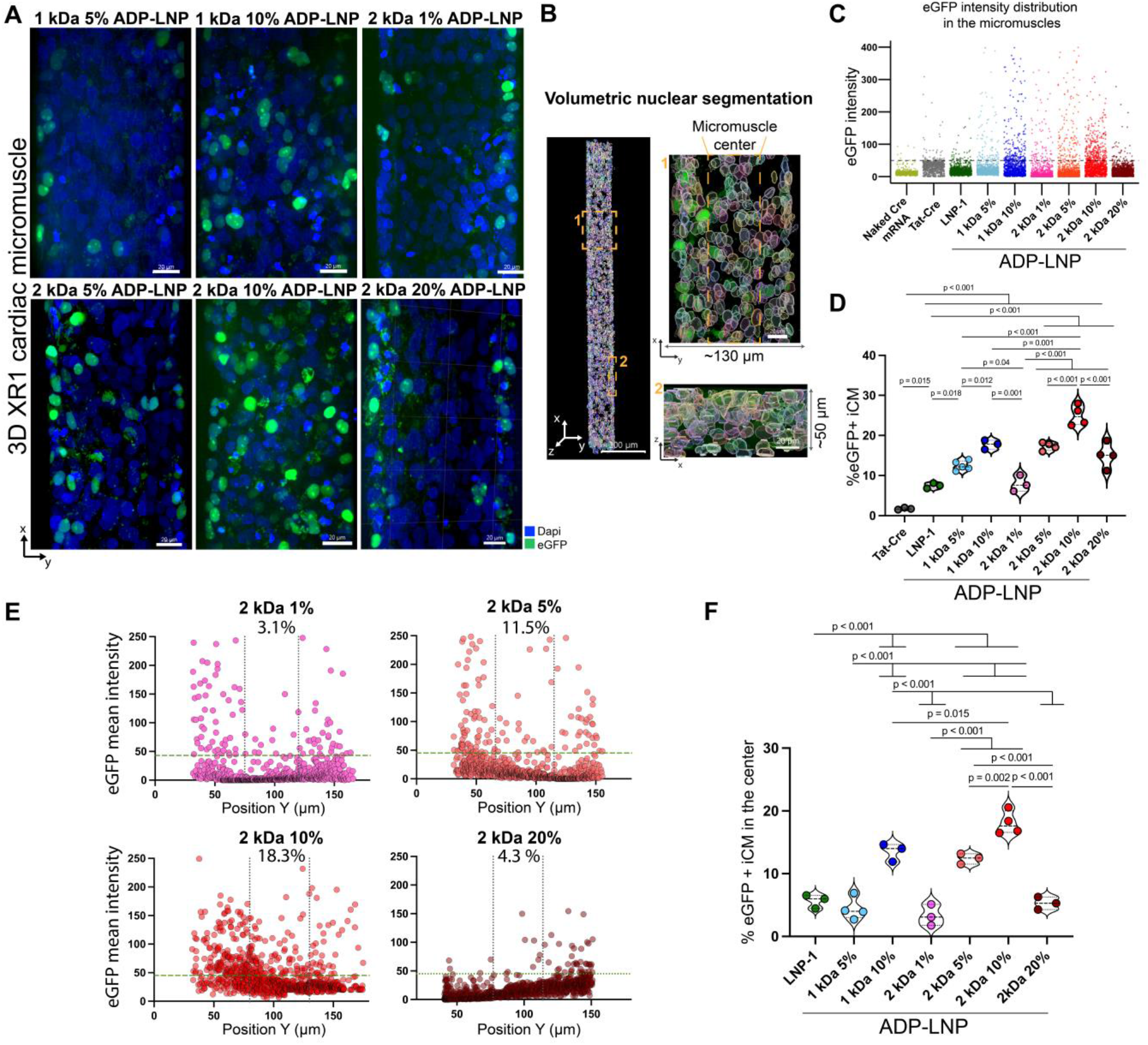
ADP-LNPs can transfect human iCMs in the cardiac MPS. **A**. Transfection (eGFP+ cells) of XR1-iCMs in the MPS exposed to the different LNP formulations. **B**. Volumetric nuclear segmentation of the cardiac micromuscle in the XY and ZX plane (inset). **C**. Representative dot plot showing the intensity distribution of eGFP+ cells after LNP/Cre mRNA delivery into the cardiac MPS. The dotted line shows the cutoff for eGFP+ cells in 3D the cardiac micromuscle. **D**. Quantitative analysis of eGFP+ cells in the cardiac micromuscle after LNP delivery of the different formulations into the cardiac MPS (n=3-5). **E**. Representative distribution of eGFP+ cells within the center of the cardiac MPS (dotted vertical lines). Each point represents an individual cell. The green line is the cutoff for eGFP+ cells. Every condition was normalized to 1100 iCMs. **F**. Quantitative analysis of E (n=3-4). Global one-way ANOVA was performed followed by post-hoc Tukey analysis.

### A higher ADP-lipid molar percentage enhanced LNP diffusion in the 3D cardiac micromuscle

To assess the combined diffusion and gene editing behavior of LNPs in dense tissues, we analyzed regiospecific LNP/Cre gene editing from the edges of the tissue chamber adjacent to the media channel to the center of the cardiac microtissues, taking advantage of the volumetric nuclear quantification described above. We quantified the percentage of eGFP+ cells in the center of the tissue based on the position of every nucleus in the cardiac micromuscle **(Fig 3a,b)** and found that microtissues treated with 2 kDa 10% ADP-LNP/Cre exhibited 18% eGFP+ iCMs in the center of the micromuscle, and this percentage significantly decreased for the other formulations **(Fig. 3e,f, Extended Data Fig. 5b)**. This result indicates that a higher ADP-lipid mole percentage in the LNPs enhanced their diffusivity, reaching the maximum for cardiac MPS where we delivered 2 kDa 10% ADP-LNP/Cre. Surprisingly, we observed a significantly lower gene editing efficiency for microtissues treated with 2kDa 20% ADP-LNP/Cre, most likely due to LNP instability or aggregation. The same trend was observed in cardiac microtissues treated with LNPs containing 1 kDa ADP-lipid whose mole percentage ranged from 5 to 10%. These results underscore the importance of considering the impact of cell-cell and cell-extracellular matrix (ECM) interactions within 3D dense tissue environments, which can significantly influence the outcomes of gene editing compared to the results previously shown in 2D monolayer screenings **(Fig. 4)**.

**Fig 4.**
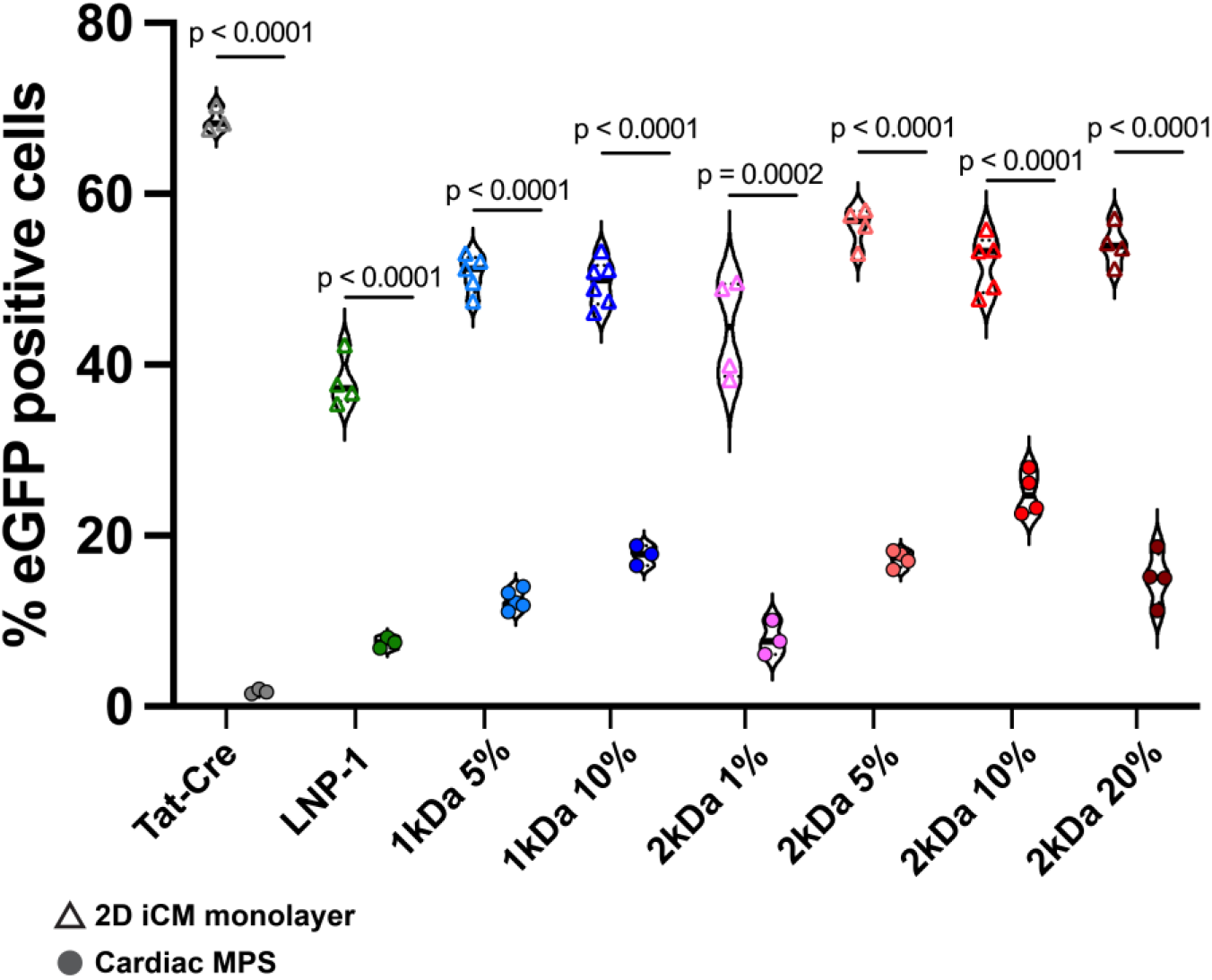
Comparison of Gene Editing Efficiency Across LNP Formulations. Gene editing efficiency of various LNP formulations are compared between 2D induced cardiomyocyte (iCM) monolayers and the Cardiac Microphysiological System (MPS). Each data point represents the observed efficiency for a specific formulation, highlighting the differences in performance across the two platforms.

### 2 kDa 10% ADP-LNP/Cre edited isogenic bi-cellular cardiac micromuscles

Cardiac fibroblasts play an essential role in the heart by maintaining and remodeling the extracellular matrix.^7, 28^ Therefore, we generated more complex micromuscles formed by isogenic XR1 hiPSC-derived cardiomyocytes and human cardiac fibroblasts (icFb) **(Extended Data Fig. 6)** with an 8:2 ratio, respectively. Four days after generating the bi-cellular cardiac micromuscles within the MPS, we delivered 2 kDa 10% ADP-LNP/Cre overnight and then kept the microtissues with flowing media for 4 more days. We observed that both cell types were successfully edited without preference for a specific cell type nor XYZ spatial location within the micromuscle **(Fig. 5a-c, Video s5)**. More importantly, the bi-cellular microtissues exhibited ∼30% eGFP+ cardiac cells upon 2 kDa 10% ADP-LNP/Cre delivery **(Fig. 5d,e)**.

**Fig. 5:**
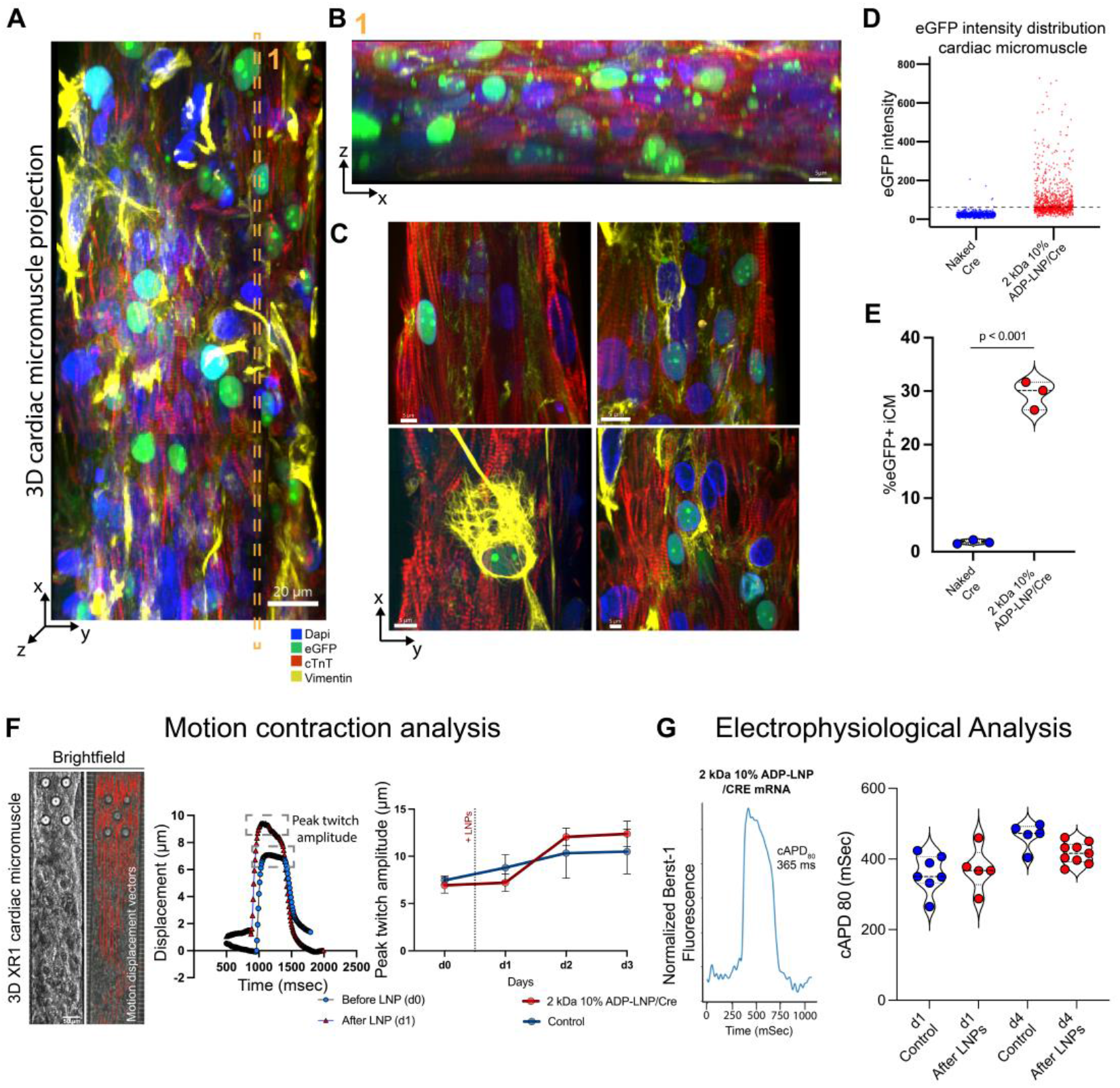
2 kDa 10% ADP-LNP/Cre edited a bi-cellular cardiac micromuscle with no toxic effects. **A**. Fluorescent image of a 3D cardiac micromuscle projection upon exposure of ADP-LNPs containing Cre mRNA. Bi-cellular cardiac micromuscle composition is 80% iCMs-20%hcFb. iCMs are identified based on the Cardiac Troponin T (cTnT) positive expression and hcFb based on the Vimentin positive expression. **B**. Section of the cardiac micromuscle exhibiting the tissue height consisting of 5-6 layers of iCMs and eGFP+ cells after LNP/Cre mRNA delivery. **C**. XY section of the cardiac micromuscle showing eGFP+ cells for iCMs and hcFb upon exposure to ADP-LNPs containing Cre mRNA. **D**. Representative dot plot showing the intensity distribution of eGFP+ cells after 2 kDA 10% ADP-LNPs/Cre delivery vs. naked Cre mRNA into the cardiac MPS. The dotted line shows the cutoff for eGFP+ cells in 3D the cardiac micromuscle. **E**. Quantitative analysis of D (n=3). Direct comparisons were made using an unpaired Student’s t-test. **F**. Motion contraction analysis over time of the cardiac micromuscle before and after LNP exposure (n=6-9). Representative brightfield images of the cardiac MPS showing the displacement vectors used to estimate the peak twitch amplitude. **G**. APD analysis over time of the cardiac micromuscle before and after LNP exposure (n=5-9).

### Electrophysiology and contractility of the cardiac micromuscle were not affected upon LNP/Cre delivery

One of the main concerns for LNPs is their clinical safety, since it is widely known that cationic lipids are inherently toxic at high concentrations and can affect the normal function of the target organ.^29, 30^ We therefore assessed the action potential duration (APD) and contractile activity of the microtissues over time for 4 days, right before (D0) and after LNP delivery (D1-3). Regarding the contractile activity, we did not observe a significant difference in the peak twitch amplitude between LNP-treated microtissues and control **(Fig. 5f, Video S6,7)**. Furthermore, the APD showed no difference compared to the controls **(Fig. 5g)**. Of note, the peak twitch amplitude and the APD exhibited a slight increase over time in both conditions due to the ongoing remodeling of the micromuscles over time. Taken together, the cardiac MPS demonstrated that the 2 kDa 10% ADP-LNP formulation is safe and effective for mRNA delivery to cardiomyocytes *in vitro*.

### *In vivo* administration of 2 kDa 10% ADP-LNP/Luc in non-infarcted mouse hearts showed increased accumulation in the heart and lower off target levels in the liver

Based on the best-performing ADP-LNP formulation identified in the cardiac MPS screening, we tested its effectiveness for *in vivo* mRNA delivery by direct intracardiac injection of 2 kDa 10% ADP-LNP and the standard LNP-1 containing firefly luciferase mRNA **(Luc) (Fig. 6a, Extended Data. Fig. 7a)**. Analysis of luciferase activity in the heart and liver showed that 2 kDa 10% ADP-LNP/Luc exhibited a significant fivefold increase in luciferase activity in the heart, in line with the results shown in the cardiac MPS. Notably, liver uptake was markedly lower compared to the standard LNP-1 formulation. Quantification of luciferase activity in isolated organs showed that the 2 kDa 10% ADP-LNP achieved approximately ninefold enrichment in the heart-to-liver ratio relative to LNP-1 **(Fig. 6b, Extended Data Fig. 7b,c)**. To determine whether the reduced hepatic delivery of ADP-LNPs resulted in an increased off-target accumulation in the spleen, we analyzed splenic uptake. The results showed that LNP-1 accumulated at higher levels in both the liver and spleen compared to ADP-LNP formulations **(Extended Data Fig. 8)**. Furthermore, despite reduced hepatic delivery, both ADP-LNP formulations exhibited a significantly lower spleen accumulation. Lastly, to assess potential cardiac toxicity, we performed echocardiographic measurements to evaluate the functional properties of mouse hearts 1 month after intracardiac delivery of 2 kDa 10% ADP-LNP/Cre mRNA **(Fig. 6c)**. No significant differences in cardiac function were observed between LNP-treated and control groups. Specifically, measurements of ejection fraction (EF) and systolic/diastolic volumes revealed no significant changes, indicating that the ADP-LNP formulation does not impair cardiac function. Collectively, these results highlight the crucial role of ADP-lipid in controlling LNP biodistribution and demonstrate the potential of 2 kDa 10% ADP-LNP to enhance cardiac-specific delivery while minimizing off-target accumulation.

**Fig. 6:**
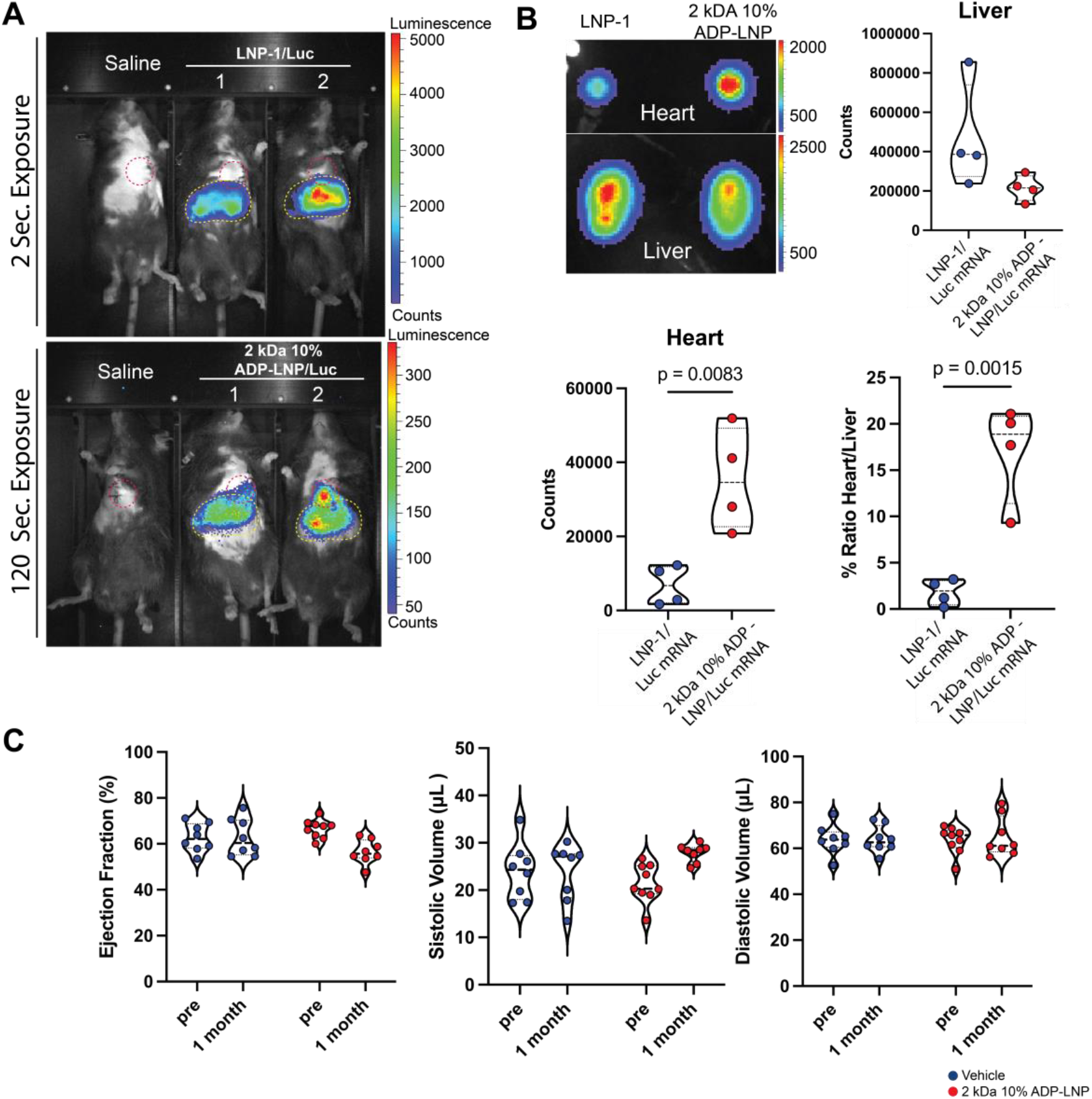
*In vivo* distribution of 2 kDa 10% ADP-LNP shows higher cardiac targeting efficiency and lower liver off target delivery when compared to LNP-1. **A**. *In vivo* Luciferase activity in anesthetized mice shows decreased liver uptake (yellow dotted circle) of 2 kDa 10% ADP-LNP when compared to LNP-1 with a concomitant increase in cardiac uptake (red dotted circle) after direct intracardiac injection. **B**. Imaging and quantification of luciferase activity in isolated heart and liver confirms that 2 kDa 10% ADP-LNP displays a much higher cardiac specificity and lower liver activity when compared to LNP-1 (n=4). Direct comparisons were made using an unpaired Student’s t-test. **C**. Echocardiographic assessment of mouse hearts 1 month after intracardiac delivery of 2 kDa 10% ADP-LNP/Luc mRNA, including measurements of Ejection Fraction (EF), Systolic Volume (SV), and Diastolic Volume (DV). No significant differences were found.

### Effective 2 kDa 10% ADP-LNP delivery to cardiomyocytes of infarcted and non-infarcted mouse hearts

To explore the functional delivery of mRNA by ADP-LNPs and identify the cells targeted in the mouse heart *in vivo*, we pursued a similar approach to the one we used for the cardiac MPS screening using the XR1 hiPSC reporter line **(Extended Data. Fig. 7a)**. For these experiments, we used Ai6 mice, which harbor a silent GFP variant (ZsGreen1) cassette that becomes activated upon Cre-mediated deletion of the stop cassette. Cre mRNA delivered via the 2 kDa 10% ADP-LNP was injected into non-infarcted control and infarcted mouse hearts, and ZsGreen1 expression was analyzed after one week of injection. High levels of ZsGreen1 signal were evident in the heart, with the strongest expression detected in the apex of the left ventricle **(Fig. 7a,b, Video S8-S12, Extended Data. Fig. 9)**. We observed the same level of ZsGreen1 expression in infarcted and non-infarcted Ai6 mouse hearts, suggesting that this LNP formulation would be equally efficient for developing new therapeutics in both cases **(Video S8-S12)**. These results indicate effective delivery of Cre recombinase *in vivo* using 2 kDa 10% ADP-LNP. To confirm these findings and to get a better understanding of the specificity of the ADP-LNP/Cre delivery, we performed an immunofluorescence staining with different cell-specific markers to identify ZsGreen1-expressing cell types in the heart. We observed that most of the ZsGreen1+ cells were identified as cardiomyocytes, as indicated by the presence of cardiac troponin I (cTnI – white asterisks) and distinctive sarcomere bands. We also observed a partial co-localization with smooth muscle and pericytes cells (αSMA+ - yellow arrows) but only rare endothelial cell transfection (IB4+ - white arrows) **(Fig. 7C)**. In summary, we found an effective LNP for mRNA delivery into the heart, where we observed a significant number of edited cardiomyocytes as well as some non-cardiomyocyte cells, although the direct injection approach precluded reliable quantification. Furthermore, the *in vivo* experiment allowed us to demonstrate that the cardiac MPS is a useful preclinical model to predict which formulations will be effective and safe for the heart.

**Fig. 7:**
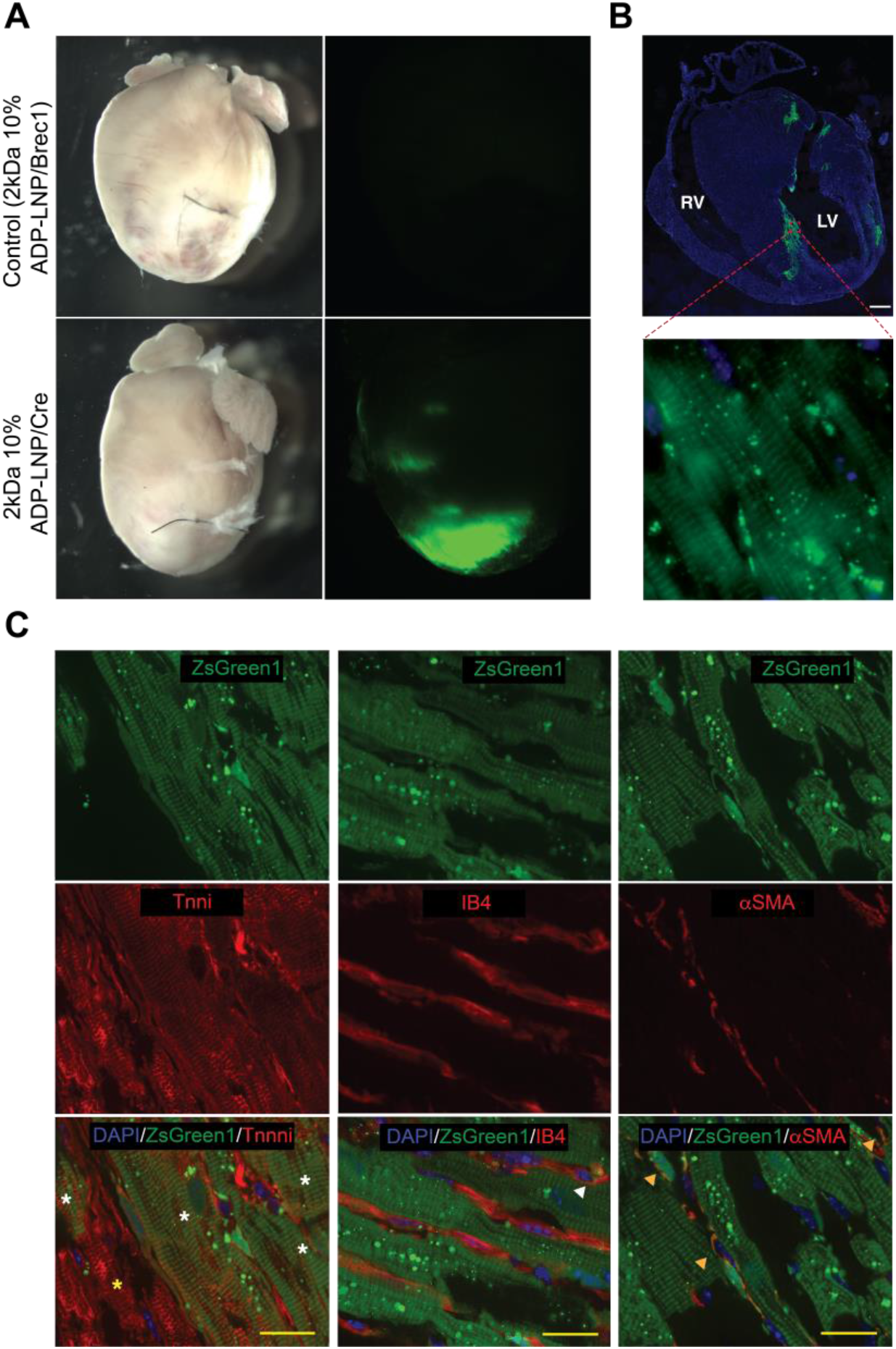
High efficiency of cardiac delivery upon intracardiac injection of 2kDa 10% ADP-LNP. (A) Brightfield (left) and fluorescence (right) whole-mount images showing ZsGreen1 activity in the apex upon intracardiac delivery of 2 kDa 10% ADP-LNP/Cre in hearts after coronary ligation. (B) Cryosections and immunofluorescence show the spatial distribution of ZsGreen1 activity in the left ventricle of the heart in homeostasis induced by Cre recombination of the stop cassette. Inset shows high magnification of cardiac ZsGreen1 positive cells with evident sarcomeres indicative of cardiomyocytes compared to large area negative for ZsGreen1. (C) Immunofluorescence staining for: (1) Cardiac cells (*Tnni*, left) – white arrows indicate colocalization with the ZsGreen1 signal, while yellow arrows highlight Tnni+ cells without colocalization; (2) Endothelial cells (*IB4*, middle) – white arrows indicate colocalization with ZsGreen1+ cells; and (3) Smooth muscle/pericyte cells (*αSMA*, right) – green arrows indicate colocalization with ZsGreen1+ cells. Scale bar: 20 μm.

## Discussion

In this study, we identified a new class of LNPs with an enhanced ability to diffuse within the heart, transfect cardiomyocytes, and promote gene editing upon LNP/Cre mRNA delivery. To this end, we used a previously developed cardiac MPS as a screening platform for various LNP formulations. Furthermore, we demonstrated that our findings in the cardiac MPS were rigorous and reproducible. We confirmed the results by testing *in vivo* the organ distribution of the most efficient ADP-LNP formulation and comparing it to a standard formulation with a non-degradable PEG-lipid conjugate. Therefore, we were able to replace, reduce, and refine the use of animal models (also referred to as the 3Rs^31^) in line with the increasing demand by society and governments to find alternatives to animal testing.^6^ This novel screening system also highlights a possible way to generate original phenotypic tissue-level approaches that may accelerate the development of safe and effective heart failure therapeutics, while significantly decreasing healthcare costs compared to current strategies.^4, 5^

We also demonstrated that the LNP screening in 2D cardiomyocyte monolayers failed to predict the results shown in the MPS, underlying how engineered tissues have the potential to replace traditional 2D models as preclinical testing platforms. One of the main reasons behind this disparity is the self-organizing tissue structure of tissue-level approaches, as the cell-cell and cell-ECM interactions have a significant influence on the maturation of cardiac tissues and interactions with different macromolecules, such as LNPs, that diffuse within them.^7, 28^ We reported previously the superiority of the cardiac MPS, particularly in regard to contractility and inotropic drug response. The MPS offers a more adult-like human hiPSC-CMs with stable electrophysiology properties and prediction of false positive and negative drug response, while 2D cultures do not.^32^ Furthermore, the incorporation of isogenic hiPSC-derived cardiomyocytes and cardiac fibroblasts in the same 3D micromuscle performed in this study allows for a better understanding of the LNP delivery in more complex microtissues. An additional advantage over 2D cultures relies on the fact that cardiac tissues have been successfully supported by mixing cell-type-specific culture media in ratios proportional to the respective cell numbers. Nevertheless, it is important to mention that the impact of suboptimal media on the function of complex tissues remains poorly understood.^7^

We also showed that the ADP-lipids allowed us to incorporate a higher level of PEGylation into the LNPs without significantly impacting mRNA delivery. This resulted in a higher LNP diffusivity within the cardiac micromuscle, due to the use of 2 kDa molecular weight, as well as the increase in the ADP-lipid mole percentage. The recent work describing the acid degradable linker development demonstrated that the enhanced efficiency results from rapid hydrolyzation in the acidic environment of early and late endosomes, consequently inducing endosomal disruption, which is in line with our results in 2D cardiomyocyte monolayers and 3D cardiac micromuscles.^18^ Collectively, we confirmed that this novel acid degradable linker based on an azide-acetal solves the stability problems associated with developing rapidly hydrolyzing linkages, resulting in a breakthrough in tackling the “PEG dilemma”.^13, 14^

Our work showed that the 2 kDa 10% ADP-LNPs outperformed the standard composition in terms of its accumulation in the heart with remarkably lower off-target levels in the liver. Taking into consideration that this administration route is more invasive and might limit clinical translation, further optimization of ADP-LNP delivery via I.V. injection or other alternatives is expected.^33^ However, functional studies upon LNP/Cre delivery demonstrated a high degree of cardiomyocyte targeting, and we observed a very similar ZsGreen1 expression profile in non-infarcted control and infarcted animals. This finding highlights the potential of this new class of ADP-LNPs in the development of either cardioprotective therapeutics or cardiac genetic disease repair.

It is worth mentioning that the majority of recent cardiac therapeutics for post-cardiac infarction treatment are focused on the re-entry of pre-existing cardiomyocytes into the cell cycle to substitute lost cells^34, 35^, reprogramming of fibroblasts to transdifferentiate into cardiomyocytes^36^, and growth factor delivery to improve vascularization and inhibition of inflammation.^37, 38^ Therefore, the development of novel non-viral carriers such as the ADP-LNPs is poised to make a significant breakthrough in cardiovascular medicine. Its higher diffusivity in tissues along with the transient expression of mRNAs will enhance transfection efficiency and minimize the potential risks of prolonged protein expression in comparison to viral vectors, such as AAV and lentivirus.^11, 39^ It is also important to highlight the lower immunogenicity triggered by non-viral vectors, as well as the lower cost of production and ease of manufacturing compared to viral carriers. Lastly, LNPs are currently the most advanced non-viral vehicle for genome editing cargo delivery *in vivo* while the size of specific genes such as the ones involved in the gene editing process can exceed the cargo capacity of commonly used virus-based gene delivery strategies.^10^

In summary, we identified an acid degradable LNP formulation with improved diffusivity and functionality upon Cre mRNA delivery by using a novel strategy to perform the screening in a cardiac microfluidic device, while reducing the need for animal models over the entirety of the characterization process.

## Methods

### LNP chemistry and characterization

The LNPs are comprised of 5 main lipids: 1) the ionizable lipid (10Z,13Z)-1-(9Z,12Z)-9,12-octadecadien-1-yl-10,13-nonadecadien-1-yl ester (**D-Lin-MC3-DMG**); 2) the cationic lipid *N*-[1-(2,3-dioleoyloxy)propyl]-*N,N,N*-trimethylammonium methyl (**DOTAP**); 3) 1,2-dioleoyl-sn-glycero-3-phosphoethanolamine (**DOPE**); 4) cholesterol; and, 5) the acid-degradable PEG-cholesterol (**1**) (**Fig. 1**). The benefits of using acid cleavable PEG chains in LNPs is widely known. However synthesizing acid-degradable lipids has been challenging due to their instability. The azido acetal linkage shown in **Figure 1** is designed to overcome the challenges associated with making acid-degradable lipids, due to its two-step mechanism of degradation. The azido acetal is a very stable acetal, due to the weak electron donating character of its azide (hammett sigma value = 0.1). This high stability allowed synthesis of the acid-degradable PEG-cholesterol (**1**) on a multigram scale. We synthesized two variants of PEG-cholesterol, which contained either a 1kDa PEG or a 2kDa PEG. Lipids stock solutions of DOTAP, D-Lin, DOPE, Cholesterol, and DMG-PEG2K at a concentration of 10 mg/mL were separately dissolved in ethanol. Commercially available DOTAP is in a solution of chloroform (25 mg/mL), and the solvent was removed under reduced pressure before use. It was then dissolved in the ethanol for storage. A stock solution of ADP (**1**) at 40 mg/mL was dissolved in DMSO. All stock solutions were stored at -30 °C. Before LNP formation, lipids were allowed to warm to room temperature or heated up to 37 °C to ensure complete dissolution, and vortexed whenever necessary. Lipids in the stock solutions were added to a 200 μL PCR tubes, following the details in tables 1 and 2, to generate a 10 μL solution of lipids. 1 μL of the mixed lipids solution was mixed with 1 μL of eGFP mRNA or Cre mRNA (1 mg/mL) and pipetted for mixing. For the ADP-LNPs, 2 μL of DTT (10 mM in PBS pH 7.4) was added to the mixture and mixed again by pipetting. The LNP solutions were incubated at room temperature for 15 min.

### Development of XR1 WTC human iPSC line

WTC-hNIL transgenic iPSC^40, 41^ were further modified by integrating the 8.4kb XR1 (e**X**cision **R**eporter 1) transgene into the CLYBL safe harbor locus. The XR1 donor plasmid was derived from pC13N-dCas9-BFP-KRAB^42^ and was a gift from Martin Kampmann by replacing the dCas9-BFP-KRAB coding sequence with a lox-STOP-lox-eGFP-P2A-mTagRFP coding sequence. The construct is designed such that CAG driven expression of the fluorescent reporter is prevented until the STOP terminator sequence is removed, either by Cre-Lox recombination or nuclease excision, similar to the Ai9 murine reporter^27^. To generate the cell line, 1 × 10^6^ WTC-hNIL iPSCs were added to 100 μL P3 buffer with 6 µg of the XR1 donor plasmid, and 3 µg of each paired TALEN (Addgene, 62196/62197) targeting the CLYBL locus (Cerbini et al., 2015). Cells were transfected using the P3 Primary Cell 4D-Nucleofector X Kit L (Lonza, V4XP-3024) with pulse code DS138. After nucleofection, cells incubated for 5 minutes at room temperature and were then plated in Stemfit media with 10 µm Y-27632 in a serial dilution into 5 wells of a 6-well plate. Non-transfected cells were plated in the sixth well. After two days, post-nucleofection media was exchanged for Stemfit containing G418 and 10 µM Y-27632. Cells were kept on G418 selection until all cells in the control wells were dead and only red fluorescent colonies remained in the experimental wells. Fluorescent colonies were manually picked and transferred to a new culture plate for expansion. Individual clones were screened for on-target integration via junction PCR by using a vector-specific primer paired with a flanking primer outside of the CLYBL homology arms in the donor vector **(Table 2S)**. The final clone (clone 4) was selected based on single copy integration that was confirmed via a ddPCR assay with a FAM-labeled probe targeting the Neomycin resistance gene and a HEX-labelled RPP30 reference probe (BioRad).

### hiPSC line culture

XR1 WTC-11 human induced pluripotent stem cells (**XR1 WTC-11 line**) were obtained through material transfer agreements from B. Conklin, Gladstone Institute and routinely checked for mycoplasma contamination. The XR1 WTC-11 hiPSC line harbors an eGFP gene downstream of a STOP cassette, which was integrated by nucleofection using TALEN and a donor plasmid. Cells were expanded on growth factor-reduced Matrigel-coated plates (Corning, 354248) in mTeSR1 Plus medium (Stemcell Technologies, 100-0276) that was changed daily, passaged at 80% confluency using Accutase (Thermo Fisher Scientific, A1110501), and plated at a density of 12,000 cells/cm^2^. For the first 24 h after passaging, the culture medium was supplemented with 5 μM Y-27632 dihydrochloride (Biogems, 1293823).

### Differentiation of hiPSC into cardiomyocytes

Once confluent, the XR1 hiPSC cells were differentiated into human iCMs utilizing a chemically defined cardiomyocyte differentiation protocol with some modifications.^21, 43^ Briefly, hiPSCs were treated with 6 μM CHIR99021 (Biogems, 2520691) for 2 days in RPMI 1640 (Thermo Fisher Scientific, 11875119) with B27-insulin (Thermo Fisher Scientific, A1895601). The cells were subsequently treated with Wnt inhibitor IWP4 (Biogems, 6861787) in RPMI/B27-for another 2 days. Between 5–11 days of differentiation, RPMI/B27-medium was used and changed every other day. Robust spontaneous contractile activity was typically observed on day 8-10 of differentiation, at which the medium was switched to RPMI/B27+insulin (Thermo Fisher Scientific, 17504044). Cardiomyocyte purity was characterized using flow cytometry for cardiac troponin T (cTnT).

### Isogenic differentiation of hiPSC into human cardiac fibroblasts (icFb)

We optimized the cardiac fibroblast differentiation for the XR1 hiPSC line following a previously published protocol **(Extended Data Fig. 7a)**.^44^ A confluent hiPSC monolayer was induced with 6 μM of CHIR99021 for 2 days, recovered in RPMI+B27-for 24 hr, and treated with 5 μM of IWP4 for 2 days. On day 5, hiPSC-derived cardiac progenitor cells (hiPSC-CPCs) were re-plated at a density of 20,000 cells/cm^2^ in advanced DMEM medium (Thermo Fisher Scientific, 12634028). On days 5 to 8, cells were treated with 5 μM of CHIR99021 and 2 μM of retinoic acid (Sigma-Aldrich, R2625) and recovered in advanced DMEM for 4 days. By this time, cells were differentiated into hiPSC-derived epicardial cells (**hiPSC-EPCs**). Next, these cells were re-plated and treated with 10 μM of FGF2 (PeproTech, 100-18B) and 10 μM of SB431542 (Selleck chemicals, S1067) in Fibroblast Growth Medium 3 (FGM3, Promocell, c-23025) for another 6 days. The cardiac fibroblasts were either passaged or frozen down for long-term storage. Additionally, we performed an XR1 cardiac fibroblast mRNA and protein characterization. We observed that icFbs expressed multiple fibroblast genes at the mRNA and protein level such as DDR2, CD90, Vimentin, TE-7, Periostin, Collagen Type I, and there was a significant mRNA upregulation of cardiac-specific markers like GATA4 and TBX20 **(Extended Data Fig. 7b-d)**.

### Fabrication of cardiac MPS

Microfluidic cardiac MPS were formed using an optimized version of the protocol described in our previous work.^19, 21^ Each MPS consisted of four multiplexed individual cell chambers (1,300 μm × 130 μm) connected to their cell-loading ports, two media ports, one media inlet, and one media outlet per multiplexed MPS. The media channels run adjacent to each side of the four cell chambers, separated by an array of fenestrations (2 µm × 2 µm high/width) to deliver nutrients while protecting the tissue from media flow shear stress. Two step photolithography was used to fabricate the microdevice silicon wafer mold, which contains the features of multiple MPS. The first photolithography step was 2 µm high for the fenestration pattern: the piranha cleaned wafer was put onto a headway spinner where SU8 2002 (Kayaku Advanced Materials, Westborough, MA) was poured and spun (first 500 rpm for 10 s with 100 rpm/s acceleration; then 2000 rpm for 30 s with 300 rpm/s acceleration). The wafer was baked for 1 min at 95°C, exposed at 80 mJ/cm^2^ UV exposure (Karl Suss MA6 mask aligner with i-line), and baked again for 2 min at 95°C. Non-crosslinked photoresist was developed in SU8 developer for 1 min 15 s, and the wafer was hard baked at 180°C for 30–45 min. The second layer for the cell chamber and media channel (55 μm high) could then be fabricated onto of the fenestration with a similar protocol. SU8 3050 (Kayaku Advanced Materials) was used and spun (first 500 rpm for 10 s with 100 rpm/s acceleration; then 3000 rpm for 30 s with 300 rpm/s acceleration). The wafer was baked slowly to allow for improved resolution by increasing the temperature from 30°C to 65°C at a rate of 10°C/min. The temperature was held at 65°C for 10 min before it was increased to 95°C at a rate of 5°C/3 min. Wafers were kept at 95°C for 45 min. The wafers were then allowed to cool down slowly to room temperature before being exposed to UV at 160 mJ/cm^2^. Alignment marks allowed for perfect overlay of both layers. Post exposure bake was the same as previously described, with only 2 min at 65°C and 10 min at 95°C. Finally, wafers were developed for 35 min in a shaker and sonicator before being hard baked at 180°C for 30–45 min. The cardiac MPS was formed by replica molding from the silicon wafer with polydimethylsiloxane (PDMS; Sylgard 184 kit, Dow Chemicals, Midland, MI) at a 10:1 ratio of Sylgard base to crosslinker. PDMS chips were then bonded to glass slides using oxygen plasma for 24 s, RF power at 21W, and 100% oxygen flow.

### Diffusion characterization

FRAP experiments were performed on live tissue samples in the cardiac MPS. An inverted ZEISS LSM 710 confocal microscope, with an argon-ion laser (488 nm), and a 403, 1.1 NA water objective was used. The experimental setting can be described as follows: frame size 512 3 256 pixels, zoom factor of 3, circular region of interest 30 mm in diameter, and pinhole setting 1 a.u. For this assay, we used the following probes: 1-40 kDa FITC-PEG (Nanocs, PG1-FC) and 4,70 kDa Dextran-FITC (Nanocs, DX4-70-FC). The working concentration for all the polymers was 100 μg/ml in PBS. Experiments started with 10 pre-bleach scanned images at low laser intensity, bleached at scan speed of 12.6 msec. at double zoom at 100% laser intensity, and followed by fluorescence recovery at low intensity. Data was normalized by the prebleach fluorescence intensity. All experiments were performed at room temperature.

### Heart micromuscle assembly within the MPS

We performed the LNP screening in cardiac MPS directly loaded with XR1 iCMs dissociated from the differentiation plate. Only iCM differentiations with a cTnT+ population higher than 80% were used in the study. On days 8-10, the hiPSC-CMs were dissociated with TrypLE 10x (Thermo Fisher Scientific, A1217703) and suspended in EB20 medium supplemented with 10 µM Y27632. A cell suspension with a density of 5,000 cells/3 μL was injected into the loading port of each MPS. After 3 min of centrifugation at 300g, MPS were inspected under the microscope. Chambers that were not filled with iCMs at this point were discarded. MPS were fed with 200 μL EB20 medium supplemented with 10 µM Y27632 into the inlet tip, and gravity allowed for constant flow to the outlet until equilibrium was reached. The following day and every day from then on, the medium was changed to RPMI/B27+insulin. To create isogenic iCM-icFb micromuscles in the MPS, we purified the iCMs by replating at a density of 100,000 cells/cm^2^ onto Matrigel, culturing in RPMI/B27+ without glucose (Thermo Fisher Scientific, 11879020) with 5 mM sodium Lactate (Sigma Aldrich, 71718) for 4 days. Cells were allowed to recover in RPMI/B27+ for 2 days. On days 9-10, iCMs were dissociated using TrypLE10X, and icFbs were dissociated using Accutase. An isogenic cardiac micromuscle was created by mixing 80% iCM-20% icFb EB20 and loaded as described above. On day 2 and every day after, heart micromuscles were fed with a medium composition of 80% RPMI/B27+, 20% FGM3 and 2 μM SB431542.

### LNP preparation and delivery

For the 2D iCM screening, we seeded 1.5 × 10^5^ iCMs/cm^2^ in each well of a 96-well plate (Corning, 3474). We dissociated and plated the cardiomyocytes following the same protocol previously described. The LNPs described in this manuscript were formulated using pipette mixing, following the protocols outlined in Wang et al.^45^ On day 4, different mRNA-LNP formulations were added to the cells in the plate and incubated overnight. First, 1 μL LNP + 1 μL mRNA (Trilink Biotech) + 2 μL DTT was mixed in a tube and incubated for 1 hr at room temperature. Then, we mixed this solution with 300 μL of RPMI/B27+insulin and added 100 μL of the LNP/media mixture to each of the wells. The following day, the medium was replaced with RPMI/B27+, and the cells were dissociated 2 days later under the different treatments to analyze the percent eGFP+ cells using flow cytometry. For the 3D cardiac MPS screening, the cells were loaded as described above. On day 4, we delivered the LNP/mRNA mix in RPMI/B27+ medium to the microtissues through the media inlet and let it flow overnight through the endothelial-like barrier that connects the media channel and the tissue chamber **(Extended Data Fig. 1)**. The following day, we replaced the medium with RPMI/B27+ and let the cardiac MPS recover for 4 days until we fixed them for confocal image analysis. For 3D screening in the cardiac MPS, the LNP preparation protocol was the same as the one described for 2D iCMs, except the LNP/mRNA concentration delivered to the microtissues was doubled. For controls, we treated the iCMs overnight with 60 μg/ml Tat-Cre protein (Sigma Aldrich, SCR508), and we also used Lipofectamine RNAiMAX (Thermo Fisher Scientific, 13778100).

### Evaluation of Encapsulation Efficiency

The encapsulation efficiency of mRNA in lipid nanoparticles (LNPs) was assessed using the Ribogreen assay (#R11490, Invitrogen). A 1× TE buffer was prepared by diluting a 20× TE buffer with deionized water. A 2% Triton X-100 buffer was prepared by diluting Triton X-100 with the 1× TE buffer, which was used to lyse the LNP/mRNA complexes. A 2000× Ribogreen stock solution was diluted with the 1× TE buffer. For the assay, 20 μL of LNP/mRNA complexes were lysed by mixing with the 2% Triton X-100 buffer in a 1:1 ratio, followed by incubation at room temperature for 5 minutes. Then, 190 μL of the 1× TE buffer containing Ribogreen was added to the wells of a black-wall, black-bottom 96-well plate, followed by 10 μL of either the lysed LNP/mRNA mixture (to determine the total mRNA content) or non-lysed LNP/mRNA (diluted 1:1 with 1× TE buffer). The plate was placed in a Spectra Max i3 plate reader, shaken for 1 minute, and incubated for an additional 4 minutes. Fluorescence intensity was measured according to the manufacturer’s protocol using excitation and emission wavelengths of 480 nm and 520 nm, respectively.

### General procedure for determining the size of LNPs

The size of LNPs containing mRNA were measured with a Malvin Zetasizer (Nano-ZS). For particle size measurements, 20 μL of LNP/mRNA complexes were prepared and diluted to a total volume of 100 μL with PBS at pH 7.4 and then analyzed on a Malvin Zetasizer.

### Gene expression in monolayer cultures

Total RNA was purified from tissues according to the manufacturer’s instructions using RNA Clean & Concentrator-5 (Zymo Research, R1015). Reverse transcription was performed using a High-Capacity cDNA Reverse Transcription Kit (Thermo Fisher Scientific, 4368814), following the manufacturer’s instructions. Expression profiles for genes of interest were determined by real-time PCR using primers (Life Technologies) and the iTaq Universal SYBR Green Supermix (Bio-Rad, 1725120) in a Bio-Rad CFBX384 real time system. Gene expression was normalized to the geometrical mean of glyceraldehyde-3-phosphate dehydrogenase (GAPDH) and Ribosomal Protein L37a (RPL37A) housekeeping values. The data was then log-transformed and relativized to the average of the biological replicates for XR1 hiPSC at day 0.

### Immunofluorescent staining in monolayer cultures

hiPSC-derived cells were dissociated and seeded onto Matrigel-coated ultra-low plate bottom 96-well plates (Perkin Elmer, 6055302). On days 4-5, cells were fixed using 4% paraformaldehyde for 15 min, then permeabilized and blocked with blocking buffer (0.05% Saponin and 3% normal goat serum (NGS) in PBS 1x) for 30 min. The cells were incubated overnight at 4°C with a 1:200 primary antibody dilution in blocking buffer. After washing them three times with PBS, cells were incubated with a 1:500 Alexa Fluor™ secondary antibody dilution at room temperature for 1 h, rinsed three times with PBS for 5 min, and stained with DAPI for 5 min at room temperature. Primary and secondary antibodies used for these studies are listed in Table 2S. Labeled cells were imaged using a widefield microscope (Lionheart FX, Biotek) and post-processed using Gen5 imaging analysis software (Biotek) and Image J.

### Flow cytometry

For the LNP screening with the 2D XR1 iCMs, cells were dissociated by incubation in TrypLE 1x, then resuspended in PBS. eGFP+ cell populations were quantified using the autosampler accessory for 96-well plates attached to the Attune NxT flow cytometer (Thermo Fisher Scientific). For iCM and icFb characterization, cells were fixed using 4% paraformaldehyde for 15 min, then permeabilized and washed using a Perm/Wash buffer (R&D, FC005). Cell suspensions were incubated with 1:200 primary antibodies in the same buffer for 30 min at room temperature. After washing, cells were incubated with 1:500 secondary antibodies in the Perm/Wash buffer for 30 minutes at room temperature. Stained cells were analyzed in the Attune NxT flow cytometer, and data was analyzed using the FlowJo software (Tree Star). The primary and secondary antibodies used for these studies are listed in **Table 3S**.

### Image acquisition for action potential and beating physiology studies

During imaging, cardiac MPS were maintained at 37°C on a stage equipped with a heating unit (Tokai Hit). Spontaneous recordings included 6-s voltage dye videos (BeRST-1) for action potential measurements, as well as 6-s brightfield videos for contractile activity studies. BeRST-1 dye was synthesized and purity-verified as previously described.^46^ For action potential recordings, MPS were first labeled overnight with 2 μM BeRST-1. After acquiring the videos, the post-experiment processing was performed with an in-house developed Python library that could analyze the fluorescence intensity over the time of the recording and the contractile activity from the brightfield videos. A NIKON TE300HEM microscope with a HAMAMATSU digital CMOS camera C11440/ORCA-Flash 4.0 was used, which recorded 100 FPS videos. For fluorescence imaging, a Lumencor (Beaverton, OR) SpectraX Light Engine and filtered with a QUAD filter (Semrock (IDEX), Rochester, NY) was used. BeRST-1 videos utilized the Far-red LED, 640 ms (4 × 4 binning, 10 ms). Videos were acquired using Nikon’s NIS-Elements software.

### MPS tissue isolation and immunofluorescence imaging

Tissues were flushed for 10 min with PBS at 25°C. Following this wash, 4% paraformaldehyde was added to the media channel for 15 min to fix the tissues. MPS were then washed with PBS twice for 5 min and carefully cut with a clean scalpel to open the device and expose the tissue. At this point, the PDMS component had the tissue structure attached to it. The tissues were then stained by submerging the PDMS blocks in different staining solutions: first, tissues were blocked with blocking buffer (3% NGS, 0.05% Saponin in 1X PBS) for 1hr at 4°C. The next day, they were submerged in primary antibodies at 1:100 concentration in blocking buffer for 24 hr at 4°C. Tissues were then washed twice at 25°C in blocking buffer for 2–3 hr and washed a third time at 4°C overnight. The secondary antibodies, along with 1:1000 DAPI (Abcam, ab108410), were incubated in blocking buffer for 24 h. Tissues were then washed twice at 25°C in blocking buffer for 2–3 hr and a third time at 4°C overnight before tissues were imaged. Then, the microtissues were reversibly mounted with 90% Glycerol (VWR, BDH1172) onto ultrathin microscope slides (Ibidi, 10812). Microtissues were imaged with the Opera Phenix high content screening system. All images were taken through proprietary Synchrony Optics ×63 water immersion lens. Images were acquired using Harmony software. We performed z-stacks over 60 μm with a step size of 0.5 μm.

### Spatial and quantitative eGFP expression analysis in the heart micromuscles

3D quantification of eGFP+ iCMs in the heart micromuscles was performed using the surface detection function of the IMARIS software (Bitplane). Objects were identified in the pipeline using the surface tool to segment DAPI or DRAQ5 positive nuclei (Thermo Fisher Scientific, 62251). Then, the eGFP intensity in each of the segmented objects was analyzed, which allowed the percentage of eGFP+ cells per microtissue to be calculated. Finally, based on the XY positions of each object, we quantified the percentage of eGFP+ cardiomyocytes along the width of the heart micromuscle. For the nuclear segmentation, we first defined the whole 3D heart micromuscle (1300 μm x 150 μm x 50 μm) as our Region of Interest (ROI). We applied a background subtraction with local contrast, considering that 12-13 μm is the maximal diameter of the largest sphere which fits into the object. Then, we applied the Split Touching Objects tool using the morphological split option with a seed point size of 4-5 μm. We finally filtered seed points based on a quality parameter to remove objects that would otherwise interfere with the interpretation of results. Once we obtained the nuclear segmentation, we used the image processing tool to adjust the eGFP intensity over the z-stack. First, we applied a Normalize Layer tool to adjust the brightness and contrast of individual z-slices to a uniform level. Then we finally applied a Gaussian filter to define the background and performed a baseline subtraction of this variable background.

### Delivery of firefly luciferase mRNA after myocardial infarction

Male C57Bl6/J mice, aged 8-10 weeks, were obtained from Jackson Laboratory (Stock No: 000664) and were positioned on a temperature-controlled small-animal surgical table to maintain body temperature (37°C) throughout the surgical procedure. Mice were anesthetized with 2-4% isoflurane, mechanically ventilated (Harvard Apparatus), and subjected to thoracotomy followed by direct intracardiac injection of 50 µl LNP containing 25 µg of modified RNA encoding for Luciferase was administrated at the heart apex using 31G insulin syringes. For luciferase analysis, we performed i.p. injection of 150 mL of XenoLight RediJect D-Luciferin (Perkin Elmer) 12-24 hours after intracardiac injection of LNP-RNA. Light emission from the firefly luciferase-catalyzed luciferin *in vivo* and *ex vivo* was detected using IVIS-Lumina S5 (Perkin Elmer) and quantified using total counts on selected areas (Supp. Figure X) and graphed using Prism 10. Student’s t-test analysis was performed in each sample tested (n=4).

### Delivery of Cre recombinase mRNA in infarcted and non-infarcted mouse hearts

This study was performed by intramyocardial injection of the LNPs in the apex of the adult heart in control and MI-induced mice, as described above. The MI was performed in 10 week mice by permanent ligation of the left anterior descending (LAD) artery, as described previously (REF). After surgery, control or MI animals were subjected to intramyocardial injection of 50 µl LNP containing 25 µg of modified mRNA encoding Cre recombinase or Luciferase (control) at the heart apex. One week after intervention, the expression of ZsGreen1 was visualized after optical clearing with CUBIC solution. Briefly, adult hearts were perfused with 10 ml of HBSS, 10 ml of CUBIC-P, 10 ml of 4% PFA and, after dissection, proceeded to tissue clearing. The tissue clearing steps include the modified CUBIC protocol that includes delipidation of hearts with CUBIC-L at 37°C degrees with agitation for 3 days, washing with PBS for 1 day, and RI matching with 50% CUBIC-R+(M)/H2O for 1 day, followed by CUBIC-R+(M) incubation for 3 days. Images were captured on UltraMicroscope Blaze (Miltenyi Biotech) or SmartSPIM (LifeCanvas Technologies) Lightsheet microscopes and processed with Imaris (ver 10.0.1). Samples were processed for cryosectioning and either visualized directly or subjected to immunohistochemistry using antibodies for cTnI, Isolectin GS-IB4-conjugated to Alexa Fluor 568 and, alpha Smooth Muscle Actin (Table 1S). Images were obtained using BZ-X810 fluorescent microscope (Keyence) and processed using Adobe Photoshop 2023 software (Adobe).

### Statistical Analysis

Data are shown as Mean ± SEM. Graphs were plotted on GraphPad Prism 9.5 and RStudio. Statistical analysis was performed using GraphPad Prism 9.5. Results with p values p < 0.05 were considered statistically significant. Global one-way ANOVA was performed when comparisons were made across more than two groups followed by post-hoc Tukey analysis. Two-factor repeated measures ANOVA was performed with two groups/conditions, where one was measured at different time points. Direct comparisons were made using an unpaired Student’s t-test.

## Reporting Summary

Further information on research design is available in the Nature Research Reporting Summary linked to this article.

## Data availability

The main data supporting the results in this study are available within the paper and its Supplementary Information. All raw and analyzed datasets generated during the study are available from the corresponding author on request. Source data are provided in this paper.

## Code availability

Contractile activity and action-potential waveforms of the cardiac micromuscles generated in the cardiac microphysiological systems were analyzed with in-house code. The code for the motion analysis can be found at https://computationalphysiology.github.io/mps_motion and for fluorescence analysis at https://computationalphysiology.github.io/mps.

## Acknowledgements

This work was funded in part by the California Institute for Regenerative Medicine DISC2-14045 (K.E.H., N.M., D.S.), the Rider Award from Innovative Genomics Institute-UC Berkeley (K.E.H., N.M., G.N.), the Siebel Stem Cell Institute seed grant award (K.E.H., G.N.), and Agilent Technologies Inc. (K.E.H., N.M.). We thank M. West (QB3 Cell and Tissue Analysis Facility, UC Berkeley) and the Molecular Imaging Center at UC Berkeley for assistance with image acquisition, analysis, and flow cytometry; We thank Henrik Finsberg and Samuel Wall from Simula Research Laboratory (Oslo, Norway) for their collaboration in the physiological analysis of the cardiac microtissues.

## Author contributions

G.N., M.W.C., H.H., D.S., N.M., and K.E.H. designed experimental studies, and performed data analysis. G.N., T.N., and B.S. performed *in vitro* experimental studies and iPSC cultures. M.W.C., T.N., Y.H., and S.L. performed *in vivo* experimental studies. H.H., and S.Z. generated the in-house acid degradable LNP components and performed LNP characterization studies. K.W., L.J., and B.C. designed and engineered the genetically modified XR1 WTC11 hiPSC-Cre reporter line used to generate the cardiac micromuscles.

