## Extended Data and Supplementary Figures for "A cardiac microphysiological system for screening lipid nanoparticle/mRNA complexes predicts *in vivo* efficacy for heart transfection"

**Extended Data Fig. 1**

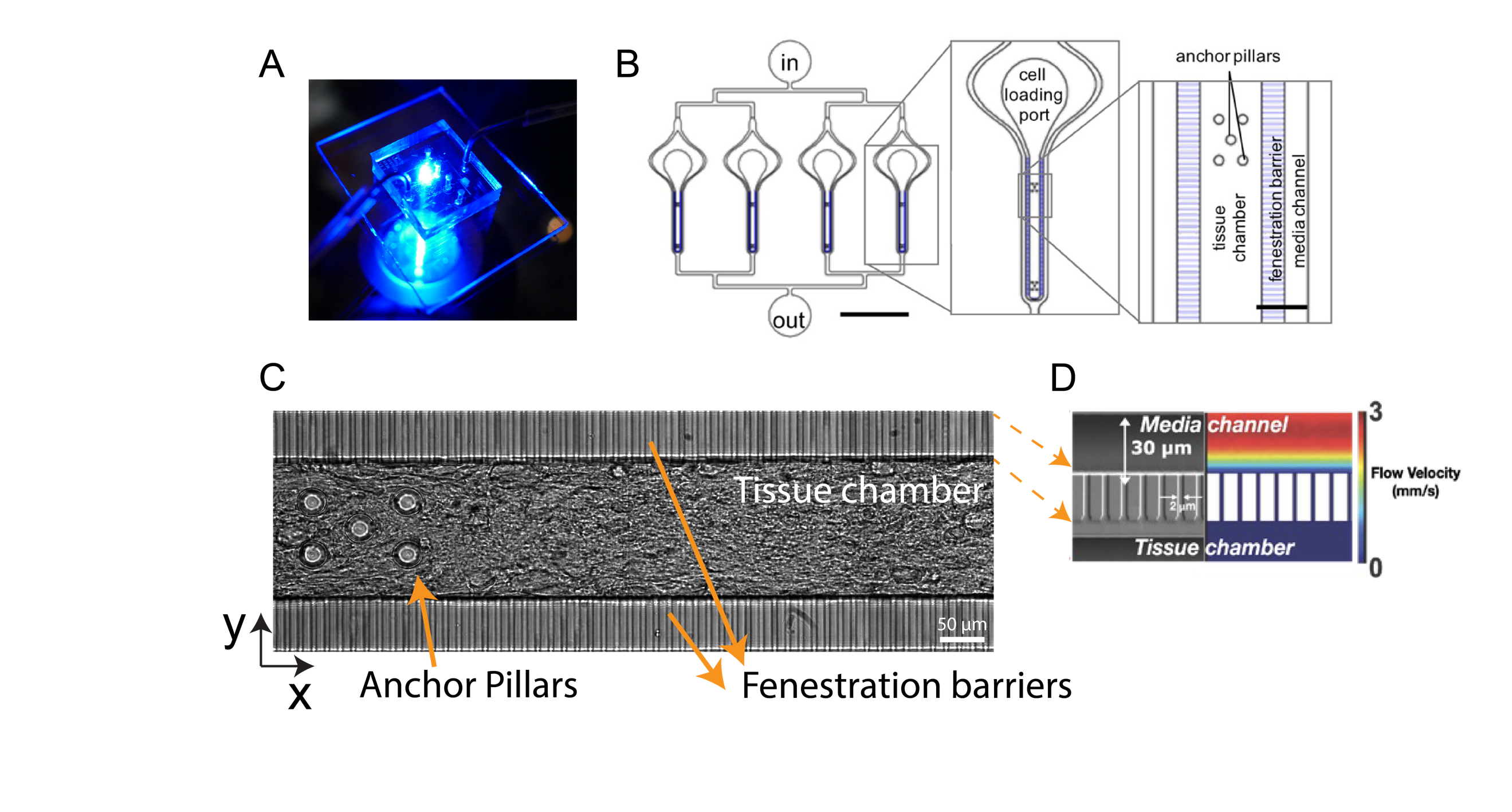

**Extended Data Fig. 1: Cardiac MPS features and function.** **A.** Photograph of a cardiac MPS under fluorescent lighting with fluidic tubing. **B.** Layout of a multiplexed cardiac MPS device comprising four parallel tissue chambers with individual cell loading ports and a common media inlet (in) and outlet (out).22 The media channels run parallel to each tissue chamber, and the fenestration barrier provides protection from fluid shear stress while allowing for the exchange of media via diffusion. Anchor pillars located at both extremities of the tissue chamber provide attachment points to keep the cardiac muscle elongated and prevent collapsing. **C.** Brightfield images of the MPS with cardiac tissues formed using XR1 WTC iCMs. **D.** SEM micrograph of the MPS showing the 2μm endothelial-like barriers connecting the nutrient channel and the tissue chamber which allows for diffusive transport of drugs and molecules. Simulated velocity profile of flow in the nutrient channel and barrier interface, demonstrating mass transport to the tissue is exclusively diffusive.

**Extended Data Fig. 2**

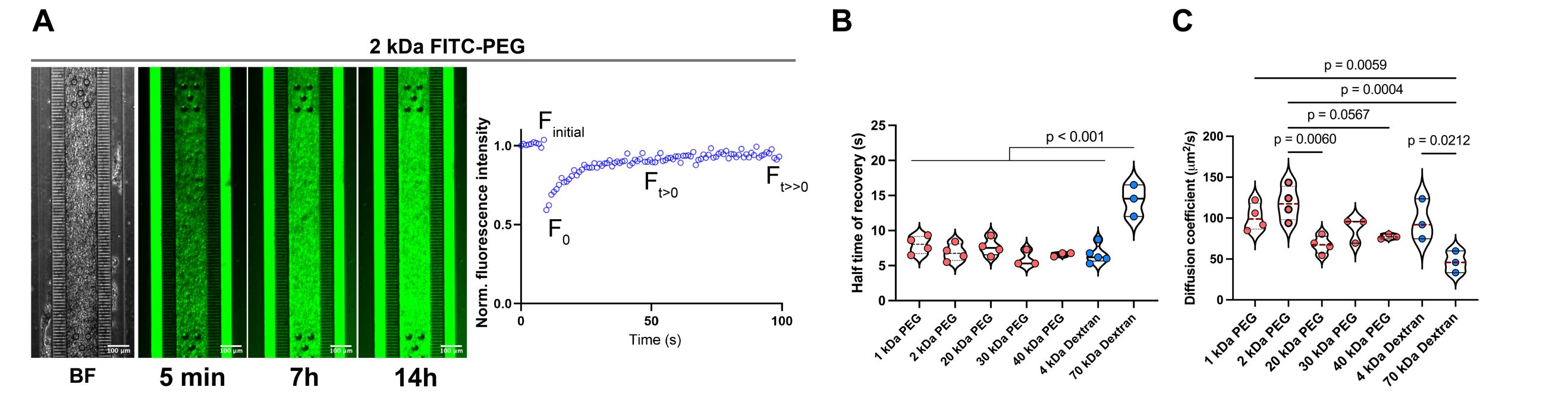

**Extended Data Fig. 2: Enhanced diffusivity of PEG polymers in the Cardiac MPS.** **A.** Time-course experiment showing the diffusion of 2 kDa PEG conjugated to FITC over time. Images were captured every 15 mins. **B.** FRAP experiment was performed after the PEG and Dextran polymers reached a steady state. Finitial is the time regime that corresponds to the initial fluorescence before bleaching; F0 is the fluorescence measurement immediately after photobleaching; Ft>0 corresponds to the recovery of fluorescence after photobleaching; Ft>>0 corresponds to the maximal recovery of fluorescence at the end of the experiment. **C.** Half-time of recovery (sec) and diffusion coefficient (μm2/s) were estimated for the different polymers in the FRAP assay.

**Extended Data Fig. 3**

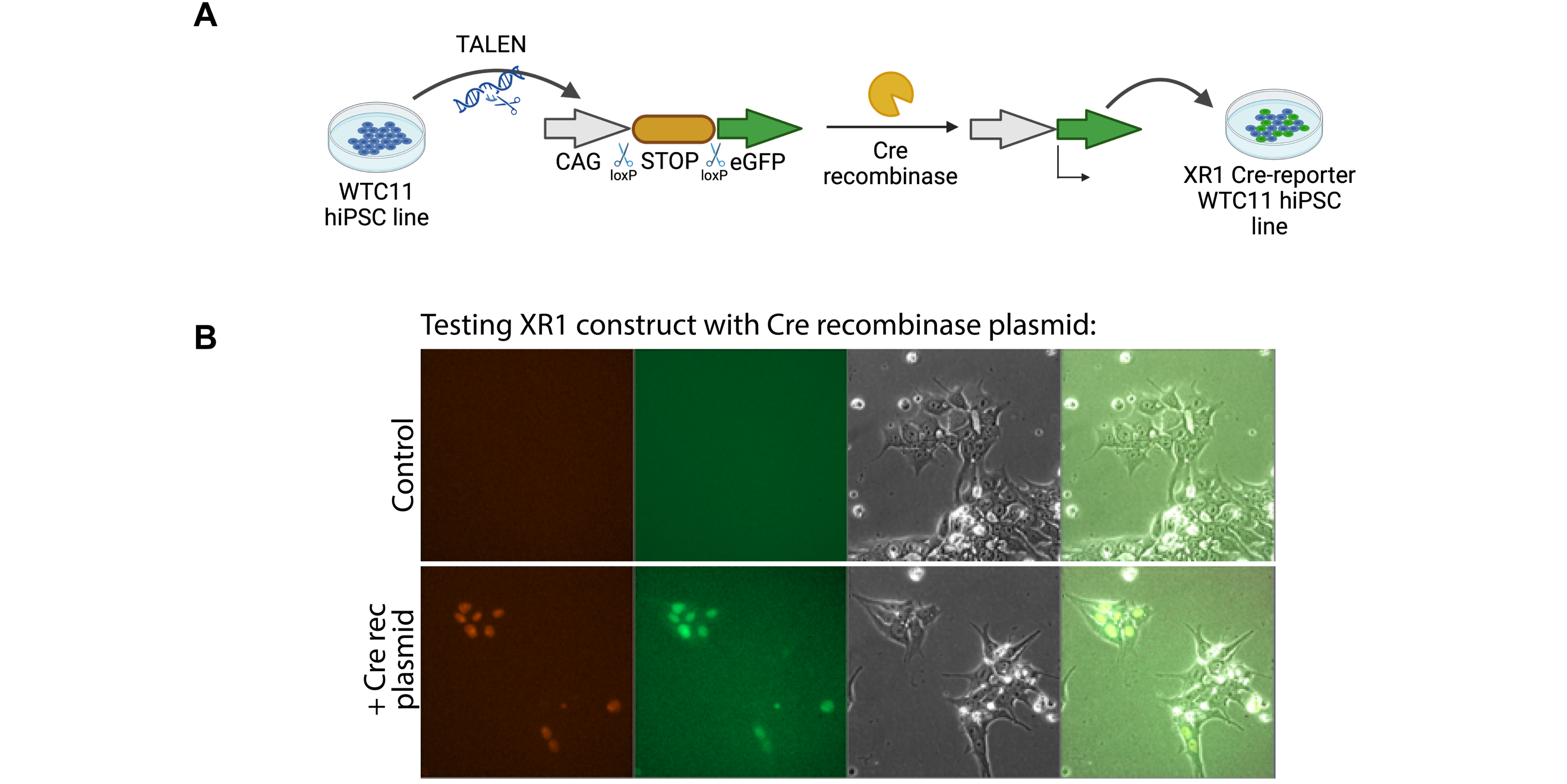

**Extended Data Fig. 3:** **XR1 WTC hiPSC Cre-reporter line development. A.** Schematic illustration of the development of a genetically modified hiPSC Cre-reporter line by TALEN-Mediated homologous recombination. **B.** eGFP expression in the XR1 hiPSC upon Cre recombinase plasmid transfection.

**Extended Data Fig. 4**

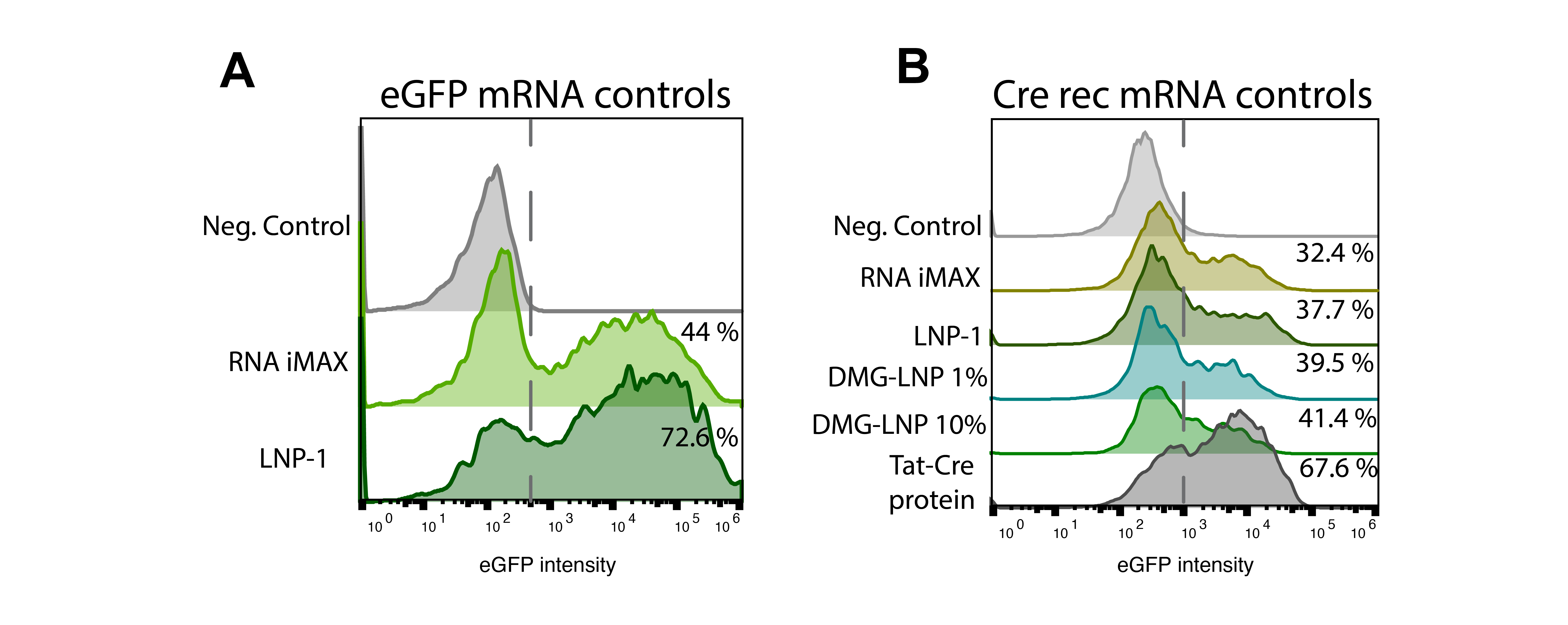

**Extended Data Fig. 4:** **ADP-LNPs can transfect human iCMs in 2D. A.** Flow cytometry analysis of 2D iCMs after RNA iMAX and LNP-1/eGFP mRNA transfection (controls). **B.** Flow cytometry analysis of 2D iCMs after RNA iMAX, LNP-1/CRE rec mRNA and Tat-Cre rec protein transfection (controls).

**Extended Data Fig. 5**

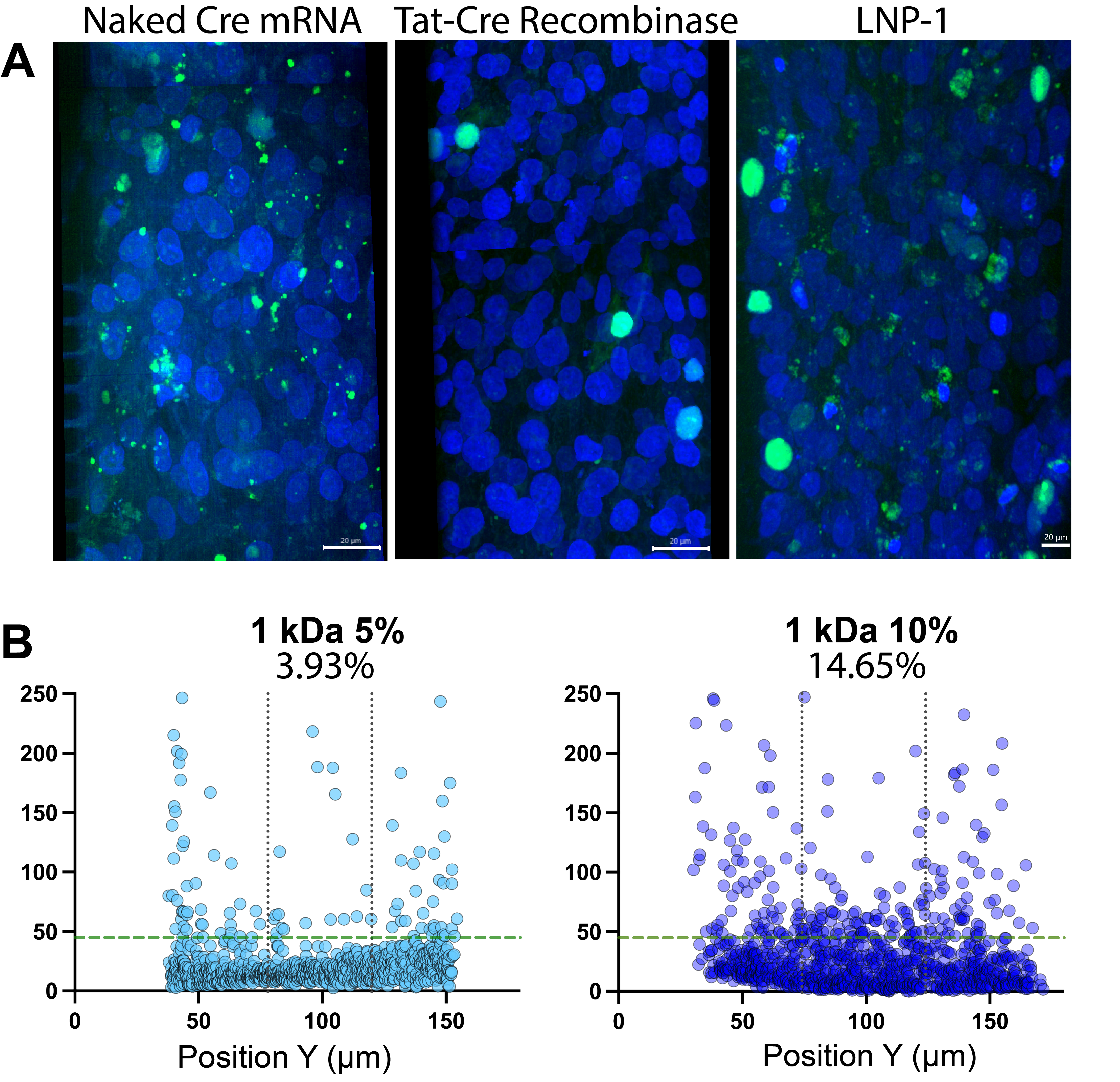

**Extended Data Fig. 5: A.** Fluorescent microscopy images showing the eGFP+ cells within the XR1 cardiac micromuscles after exposure to naked CRE rec mRNA, Tat-CRE Rec protein, and LNP-1/CRE rec mRNA. **B.** Representative distribution of eGFP+ cells within the center of the cardiac MPS (dotted vertical lines). Each point represents an individual cell. Green line is the cutoff for eGFP+ cells. Every condition was normalized to 1100 cells.

**Extended Data Fig. 6:**

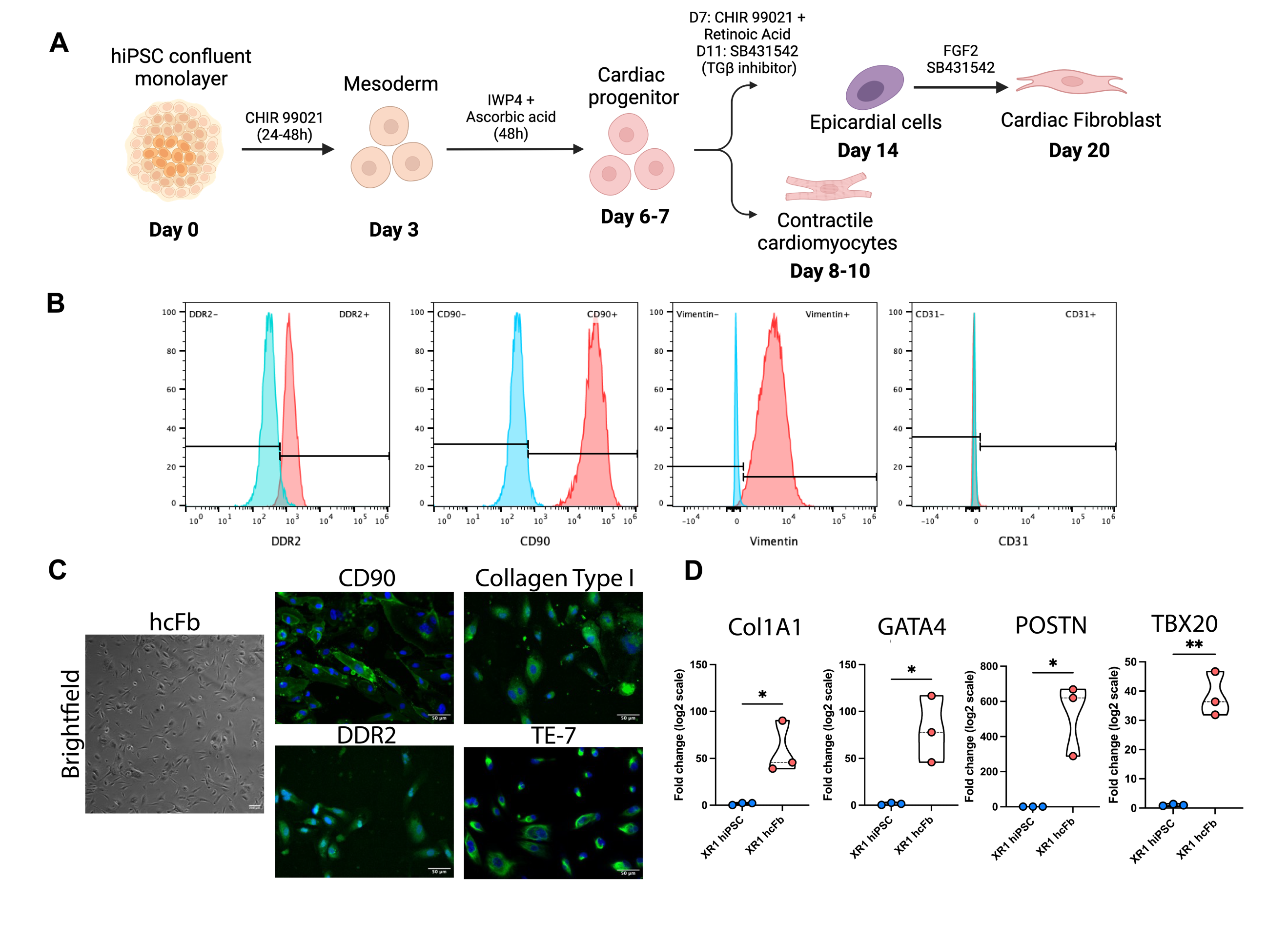

**Extended Data Fig. 6: Isogenic XR1 cardiomyocyte and cardiac fibroblast differentiation. A.** A schematic diagram showing the protocol for small molecule-directed differentiation to hiPSC-derived cardiomyocytes and cardiac fibroblasts. **B.** Representative flow cytometry analysis showing protein levels of hcFb markers such as CD90, DDR2, and Vimentin. The protein expression for the endothelial marker CD31 was also assessed. **C.** Representative brightfield image of the hcFbs in culture and immunofluorescence images showing the positive expression for hcFB markers such as Collagen Type I, CD90, DDR2, and TE-7. **D.** mRNA expression analysis of genes that are significantly expressed in cardiac fibroblasts such as COL1A1, GATA4, POSTN, and TBX20 relative to their pluripotent state. Direct comparisons were made using an unpaired Student’s t-test (*p < 0.05, **p < 0.001).

**Extended Data Fig. 7:**

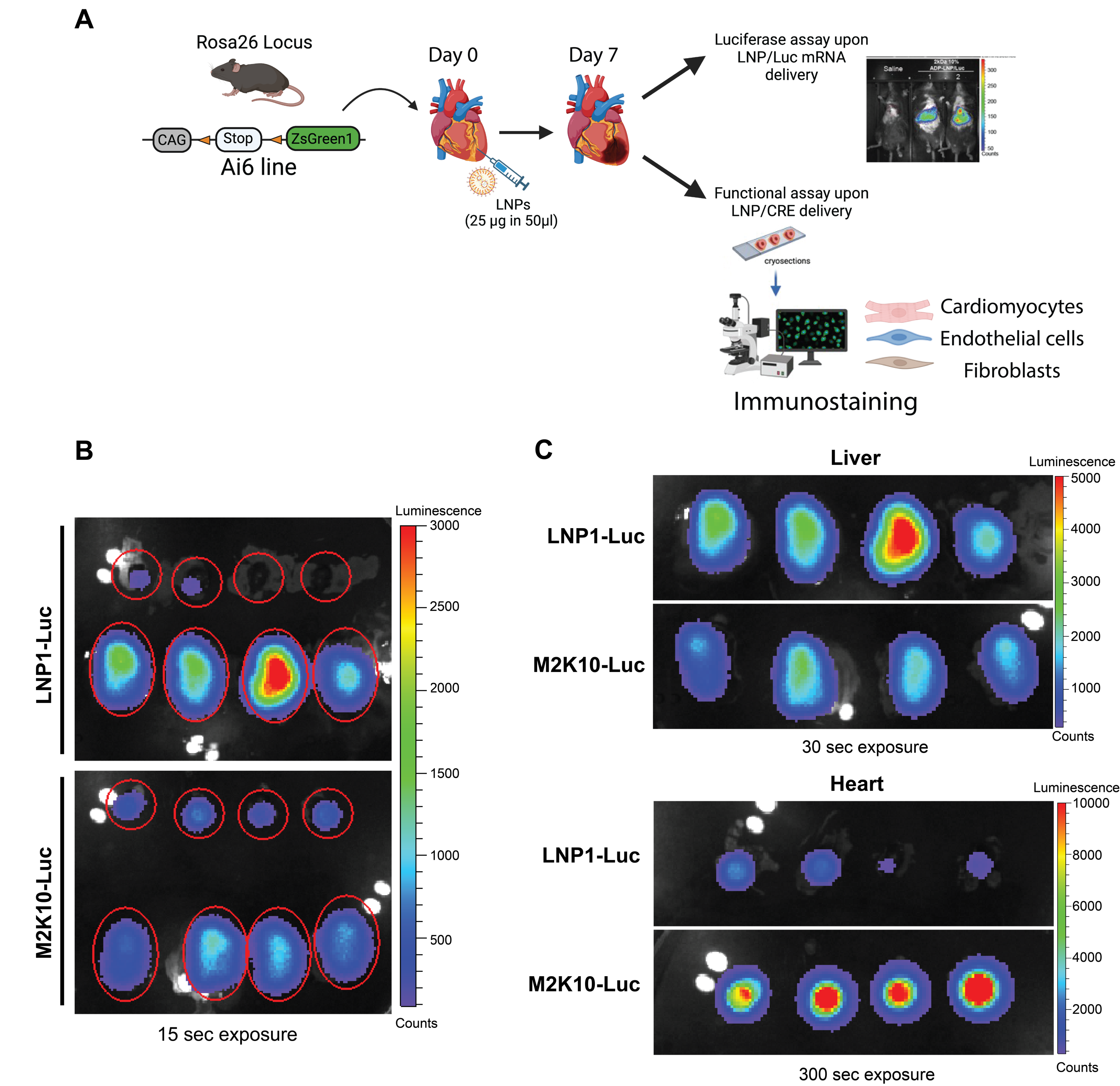

**Extended Data. Fig. 7: A.** Schematic illustration describing transfection of Ai6 mouse with either Luc mRNA or Cre mRNA delivered by 2 kDa 10% ADP-LNPs and LNP-1. **B.** Characterization of the region used in quantification presented in Fig. 5B. upon 15 seconds of exposure. **C.** Normalized images of Liver (top, exposure 30 seconds) and hearts (bottom, exposure 300 seconds) show clear increase in heart luciferase activity in 2 kDa 10% ADP-LNP when compared to LNP-1 with a small decrease in overall off-target liver expression.

**Extended Data Fig. 8:**

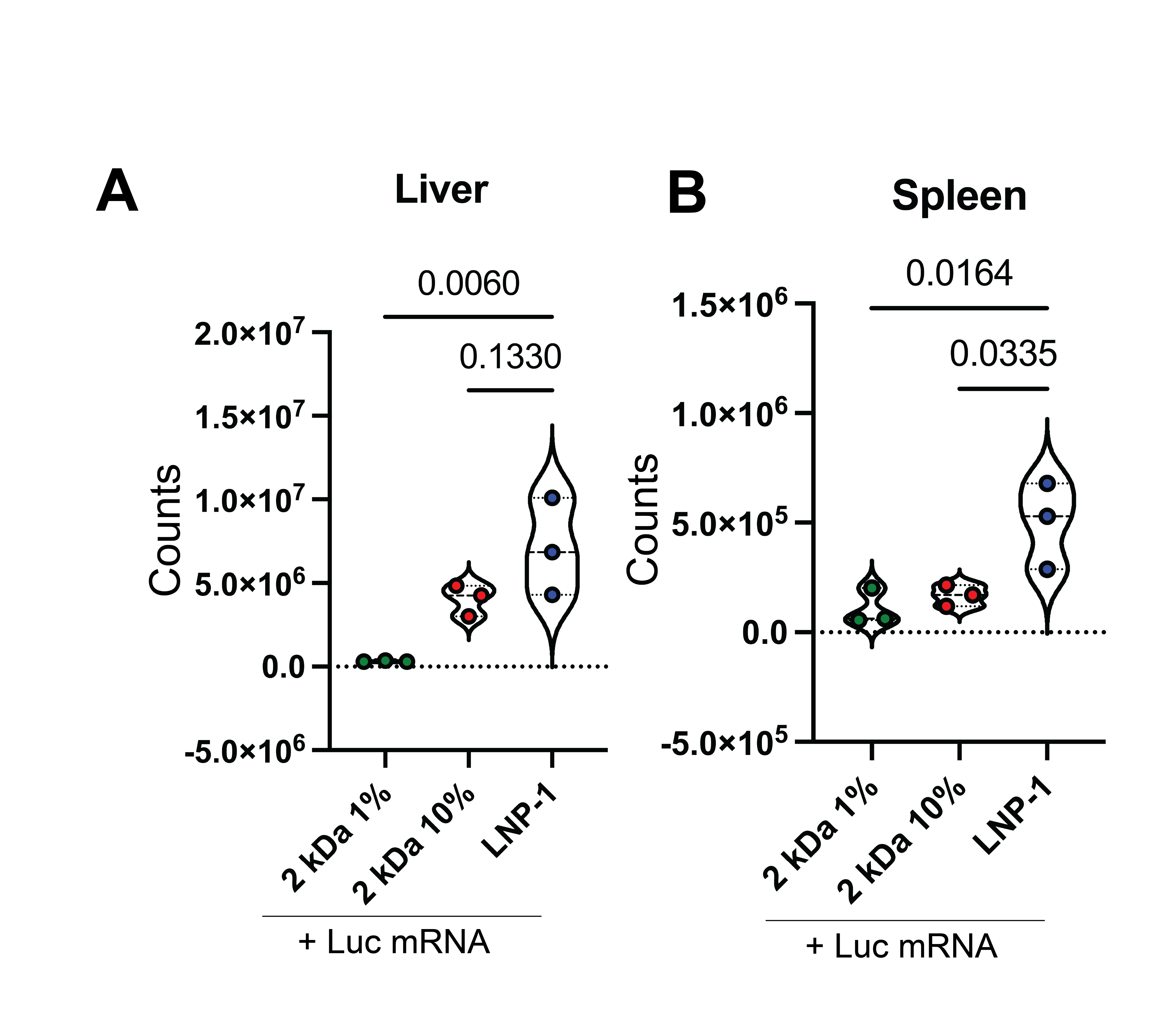

**Extended Data. Fig. 8:** Quantification of luciferase activity in isolated liver **(A)**, and spleen **(B)** following intracardiac delivery of 2 kDa ADP-LNP formulations (1% and 10% PEGylation) and the standard LNP-1 formulation (n = 3). ADP-LNP formulations show reduced off-target activity in the liver and spleen compared to LNP-1.

**Extended Data Fig. 9:**

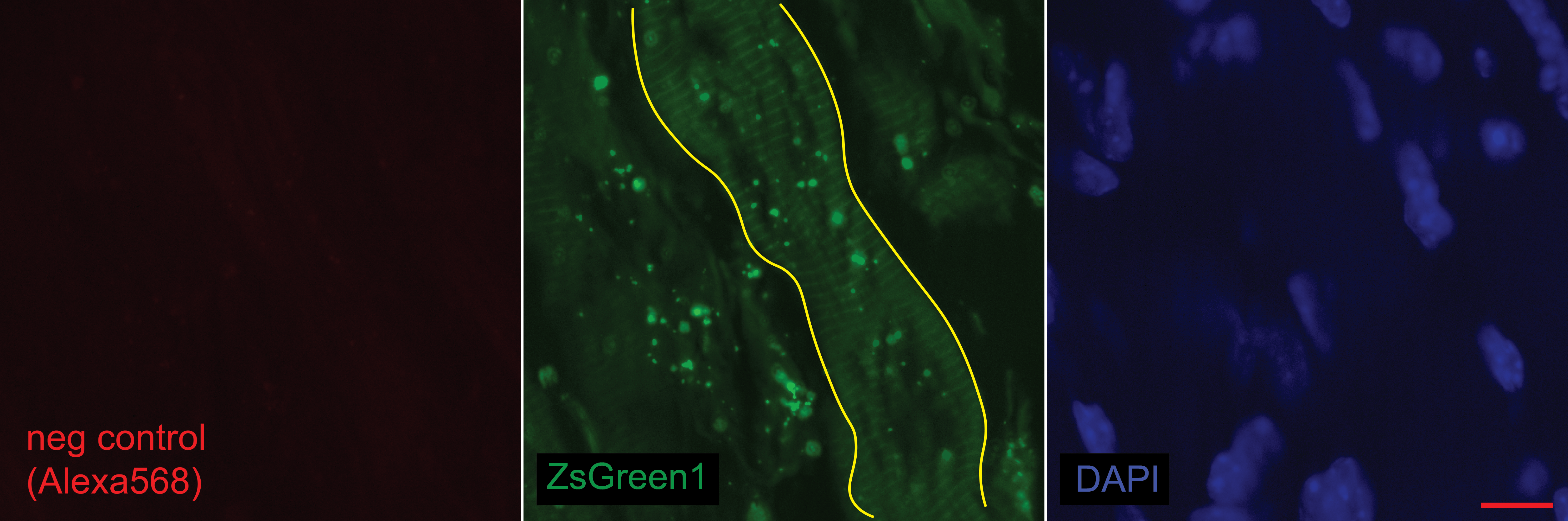

**Extended Data Fig. 9.**  Immunofluorescence control for sections shown on Fig. 7 C showing absence of signal when no primary antibody for Tnni or aSMA and no conjugated IB4 were used (neg control, left), while genetic ZsGreen1 labeling upon Cre mediated deletion shows CM with distinct sarcomeres highlighted with yellow lines (middle) and DAPI marking cell nucleus (right). Scale bars-10mM.

**Supplementary Information**

**Table 1S:** Components of ADP-LNPs with 1kDa PEG molecular weight.

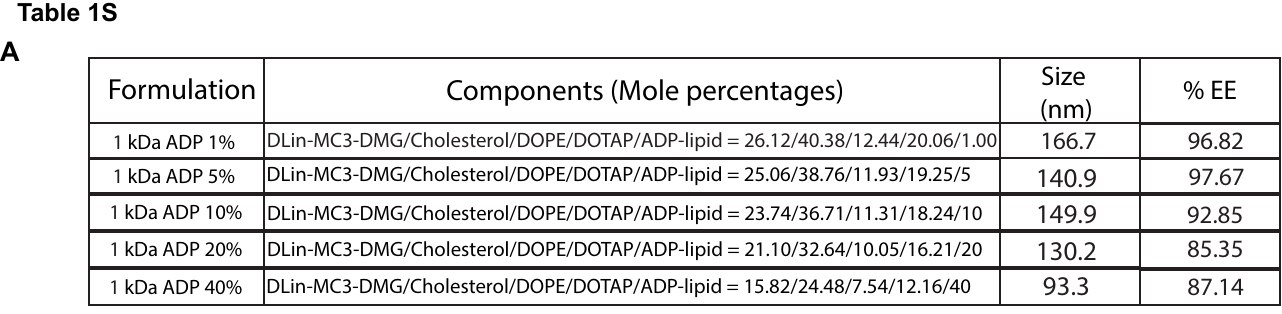

**Table 2S**

| **5’ junction primers** | **Sequence** | **Amplicon size** |
| --- | --- | --- |
| KW064 | tctagagcctctgctaaccatgttcatg | 495bp |
| KW067 | cattgatgagtttggacaaaccacaac |  |
| **3’ junction primers** | **Sequence** | **Amplicon size** |
| KW066 | aagtcgtgctgcttcatgtg | 448bp |
| KW070 | gcattctagttgtggtttgtcc |  |

**Table 3S**

| **Antibody** | **Source** | **Catalogue Number** |
| --- | --- | --- |
| Anti-CD90 | Abcam | 123511 |
| Anti-cTnT | Epredia | MS295P1 |
| Anti-DDR2 | Abcam | 63337 |
| Anti-Collagen I | Abcam | 21286 |
| Anti-TE7 | EMD Millipore | CBL271 |
| Anti-CD31 (PECAM-1) | Invitrogen | 14-0319-82 |
| Anti-Vimentin | Cell Signaling | 57415 |
| Anti-cTnI | Thermo Scientific | 701585 |
| Anti-Isolectin | Thermo Scientific | I21412 |
| Anti-alpha SMA | Abcam | Ab7817 |
| Alexa Fluor 488 goat anti-mouse | Invitrogen | A11001 |
| Alexa Fluor 647 goat anti-rabbit | Invitrogen | A21245 |
| Alexa Fluor 568 goat anti-mouse | Invitrogen | A11011 |
